# Sampling strategies and integrated reconstruction for reducing distortion and boundary slice aliasing in high-resolution 3D diffusion MRI

**DOI:** 10.1101/2023.01.11.523645

**Authors:** Ziyu Li, Karla L. Miller, Jesper L.R. Andersson, Jieying Zhang, Simin Liu, Hua Guo, Wenchuan Wu

## Abstract

**Purpose:** To develop a new method for high-fidelity, high-resolution 3D multi-slab diffusion MRI with minimal distortion and boundary slice aliasing.

**Methods:** Our method modifies 3D multi-slab imaging to integrate blip-reversed acquisitions for distortion correction and oversampling in the slice direction (kz) for reducing boundary slice aliasing. Our aim is to achieve robust acceleration to keep the scan time the same as conventional 3D multi-slab acquisitions, in which data are acquired with a single direction of blip traversal and without kz-oversampling. We employ a two-stage reconstruction. In the first stage, the blip-up/down images are respectively reconstructed and analyzed to produce a field map for each diffusion direction. In the second stage, the blip-reversed data and the field map are incorporated into a joint reconstruction to produce images that are corrected for distortion and boundary slice aliasing.

**Results:** We conducted experiments at 7T in six healthy subjects. Stage 1 reconstruction produces images from highly under-sampled data (R=7.2) with sufficient quality to provide accurate field map estimation. Stage 2 joint reconstruction substantially reduces distortion artifacts with comparable quality to fully-sampled blip-reversed results (2.4× scan time). Whole-brain in-vivo results acquired at 1.05mm isotropic resolution demonstrate improved anatomical fidelity compared to conventional 3D multi-slab imaging. Data demonstrate good reliability and reproducibility of the proposed method over multiple subjects.

**Conclusion:** The proposed acquisition and reconstruction framework provide major reductions in distortion and boundary slice aliasing for 3D multi-slab diffusion MRI without increasing the scan time. This method has the potential to provide high-quality, high-resolution diffusion MRI.

## 1. Introduction

Diffusion magnetic resonance imaging (MRI) probes tissue at the microscopic scale non-invasively^1, 2^, providing information about healthy and pathological changes to neural architecture. High-resolution diffusion MRI can depict microstructure details in the brain, facilitating tracking of thin fibers and accurate depiction of complex fiber configurations^3-6^. Because of these advantages, high-resolution diffusion MRI is compelling for neuroscientific research and clinical diagnosis.

Three-dimensional (3D) multi-slab acquisition has great potential to achieve high-resolution *in-vivo* diffusion MRI, which can produce optimal signal-to-noise ratio (SNR) efficiency for spin-echo-based diffusion MRI due to its compatibility with a short TR=1-2s and achieve thin slices using a 3D k-space encoding^7-11^. 3D multi-slab acquisitions divide the whole image volume into multiple thin slabs (typically with 10-20 slices in each slab) and encode each slab with a 3D k-space readout, typically using a 3D echo-planar imaging (EPI) trajectory for an efficient acquisition^7-9^. However, the image quality of 3D multi-slab imaging with EPI-based trajectories is compromised by two major image artifacts: distortion and boundary slice aliasing.

First, similar to conventional 2D EPI, 3D EPI suffers from distortions from the static (B0) magnetic field inhomogeneity^12^. These distortions occur along the phase-encoding direction due to its low bandwidth, especially near tissue/bone and tissue/air interfaces with large susceptibility differences. EPI distortions are conventionally corrected after image reconstruction using a field map acquired from a separate scan, using either a GRE field mapping sequence^12^ or a pair of EPI images acquired with opposite phase-encoding directions^13^. However, field mapping-based corrections are inadequate, particularly if distortions become sufficiently severe that the signal from multiple voxels overlap in the reconstructed image. Moreover, static field maps cannot capture dynamic B0 field changes due to subject motion, eddy currents, and field drift across different diffusion directions^14^.

EPI distortion corrections using a pair of blip-reversed phase-encoding images have also been developed, which have demonstrated superior performance compared to the field mapping-based correction in diffusion MRI^13, 15-19^. While most of these approaches perform distortion correction using separately reconstructed blip-up and blip-down images^13, 15, 16^, methods that jointly reconstruct blip-up and blip-down data have also been developed^17-19^. The recently proposed BUDA (blip-up/down acquisition) EPI^18^ method performed interleaved blip-reversed acquisitions for each diffusion direction and incorporated field maps estimated from separate blip-up/down images into a distortion corrected joint reconstruction. The method demonstrated improved distortion correction, dynamic B0 field mapping capability, and reduced g-factor penalty due to the combination of blip-up/down data in the joint reconstruction. However, existing blip-reversed acquisition methods require doubled scan time compared with conventional single-shot EPI using a single phase-encoding acquisition.

A second source of artifacts in 3D multi-slab imaging is slice aliasing near the slab boundary. Boundary slice aliasing arises due to the inability to achieve a sharp excitation profile, which means some tissue beyond the targeted slab is excited. The transition bands and sidelobes of the RF profile thus extend to adjacent slabs, leading to aliasing of signal from one end of the slab into the other^20^. To avoid aliasing, conventional methods expand the field of view (FOV) along the slice direction for each slab via oversampling^9^ or increase the overlapping between adjacent slabs^9^. However, these methods inevitably require additional scan time.

In this work, we aim to incorporate the blip-reversed acquisition and kz oversampling into 3D multi-slab diffusion imaging to minimize distortion and boundary slice aliasing without increasing the scan time. However, achieving this goal is fundamentally challenging. Acquiring both blip-up and blip-down images without increasing the scan time requires an extra two-fold (2x) acceleration for time compensation. The current 3D multi-slab diffusion MRI acquisition typically uses 3D multi-shot EPI with each EPI readout covering a single kz plane. Because ∼50% of sequence time is dedicated to diffusion preparation, shortening the readout with in-plane acceleration will not provide significant scan time reduction. Instead, the most effective way to reduce scan time is to accelerate along the slice direction, which determines the number of excitations required to sample along kz. This is challenging for *in-vivo* diffusion MRI because each slab needs to be sufficiently thin (i.e., 10–20mm) to ensure that the motion-induced phase variation within each slab can be accurately measured with a 2D navigator^9, 21, 22^, which leads to very limited coil sensitivity variation along the slice direction. Therefore, direct under-sampling along the slice direction will result in a highly ill-posed reconstruction. Moreover, to additionally address the boundary slice aliasing, oversampling along kz is needed, which will lead to an even higher acceleration factor and thus a more challenging reconstruction problem.

We propose an approach for 3D EPI that enables distortion- and boundary slice aliasing-corrected reconstruction similar to BUDA^18^ but with no increase in scan time compared to the conventional 3D multi-slab acquisition^7-9^ (Fig. 1). We acquire an equal number of shots with blip-up and blip-down phase encoding. The reconstruction proceeds in two stages. In the first stage, we reconstruct the highly under-sampled blip-up and blip-down images separately in order to estimate the field map^13, 23, 24^. In the second stage, we reconstruct a final image that is distortion corrected and has a more modest acceleration using all segments and accounting for the field map. In addition, we enlarge the FOV along the slice direction to reduce the boundary slice aliasing. This approach relies on the highly-accelerated reconstruction for stage 1 being sufficiently robust to accurately estimate the field map. To support this aim, we use a 3D EPI trajectory with blipped CAIPI^25^ and partial Fourier along kz. A high-SNR SPIRiT-based^26^ reconstruction is also optimized and employed. Our framework is validated with *in-vivo* experiments on a 7T scanner and achieves whole-brain diffusion imaging at 1.05 mm isotropic resolution. The diffusion analysis and tractography results reveal the higher anatomical fidelity of the proposed method compared to the conventional 3D multi-slab imaging, demonstrating the potential of our method to facilitate high-resolution diffusion MRI with improved image quality to benefit neuroscientific research.

**Figure 1.**
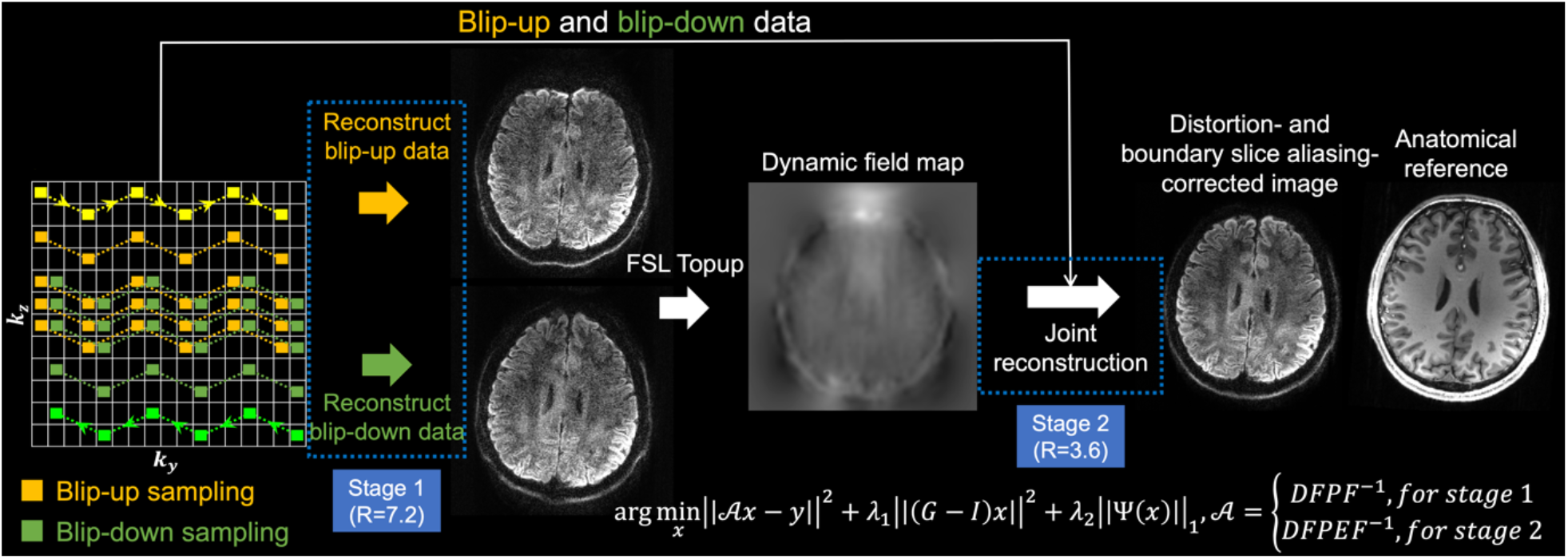
Proposed framework. The proposed sampling pattern with kz blipped-CAIPI and complementary partial Fourier for blip-up (yellow) and blip-down (green) data and the two-stage reconstruction. The trajectory of one shot of the multi-shot sampling is marked in bright yellow and bright green for blip-up and blip-down sampling with arrows indicating the phase encoding direction. The forward operator 𝒜 in the reconstruction includes Fourier Transform *F* and inverse Fourier Transform *F*^−1^, the phase modulation P representing motion-induced phase errors measured by 2D navigators, k-space sampling operation D in both stages 1 and 2, and an additional distortion operation E (captured by the field map) for stage 2. Example images are from a single volume diffusion MRI dataset (1.05 mm isotropic resolution) of a representative subject, with anatomical image listed for reference (acquired with MPRAGE at 0.86 mm isotropic resolution).

## 2. Methods

### 2.1. Blip-reversed acquisition and reconstruction for 3D multi-slab imaging

#### 2.1.1. Acquisition

Integrating 3D multi-slab diffusion MRI with blip-reversed EPI requires the acquisition of two 3D EPI images with reversed phase encodings, which can be achieved at a cost of doubled scan time. One approach to shorten the scan time is to perform under-sampling along kz. By acquiring half the segments with blip-up phase encoding and the other half with blip-down phase encoding, the total scan time of the integrated blip-reversed acquisition is identical to conventional 3D multi-slab EPI^7-9^. Importantly, 3D multi-slab diffusion MRI requires very thin slabs to facilitate the correction of motion-induced errors^8, 9, 22^. This severely limits the variation of coil profiles along the slab direction, and as a result conventional rectangular under-sampling with integer reduction along ky and kz (e.g., Fig. 2c, iii) may suffer from a high g-factor penalty and residual aliasing.

**Figure 2.**
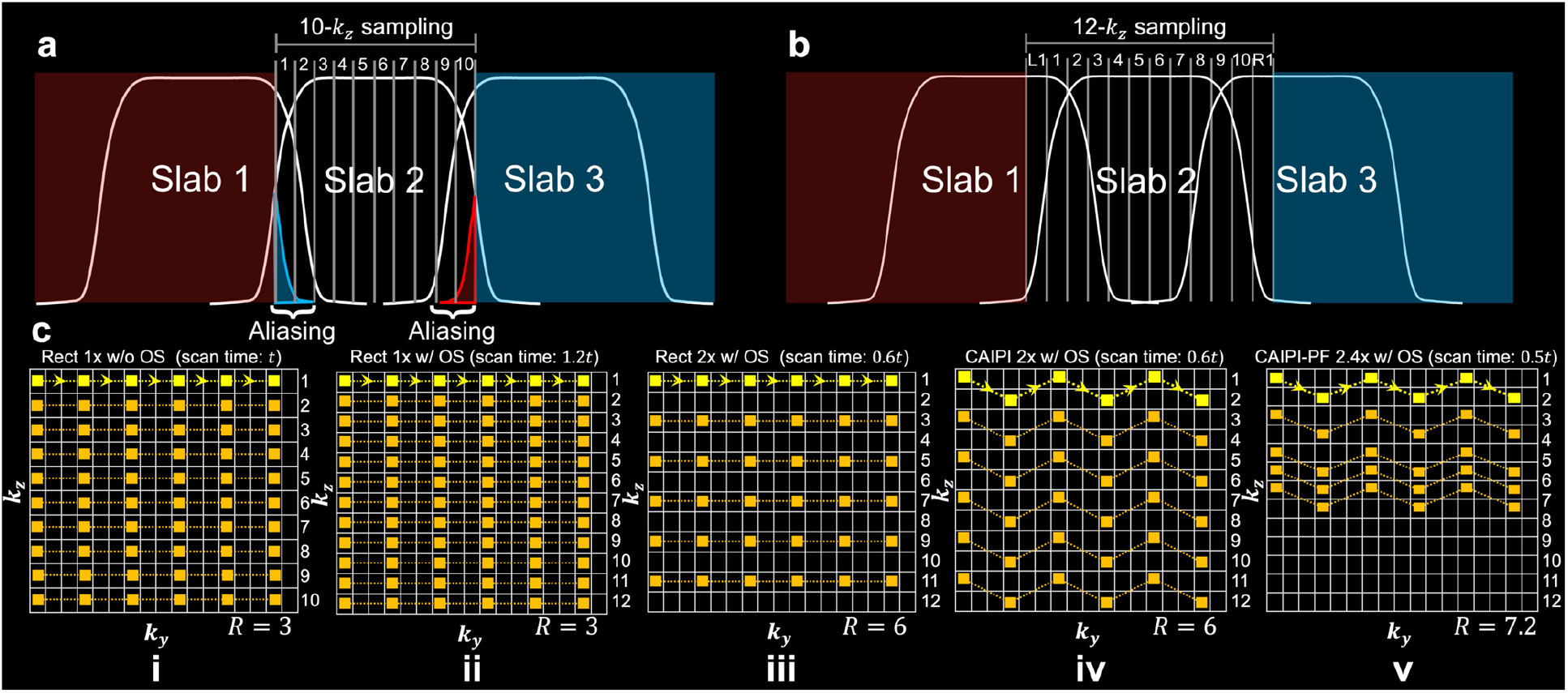
3D multi-slab acquisition and sampling patterns for blip-up data. a. Demonstration of slice aliasing at slab boundary caused by non-rectangular RF profile and limited field-of-view (FOV) along the slice direction for 3D multi-slab acquisition (10-kz sampling). b. Correction of slice aliasing by expanding the FOV for each slab through kz over-sampling (12-kz sampling) (L1 and R1 denote the 1 oversampled slice on the left and right side). c. Comparison of various sampling patterns: rectangular sampling without oversampling along kz (Rect 1x w/o OS) (i); rectangular sampling with 20% oversampling along kz (Rect 1x w/ OS) (ii); rectangular sampling with 20% oversampling and 2x acceleration along kz (Rect 2x w/ OS) (iii); CAIPI sampling with 20% oversampling and 2x acceleration along kz (CAIPI 2x w/ OS) (iv); CAIPI-PF sampling with 20% oversampling, partial Fourier and 2.4x acceleration along kz (CAIPI-PF 2.4x w/ OS) (v). A 3x acceleration along ky is applied in all sampling patterns. The trajectory of one EPI shot is marked in bright yellow with arrows indicating the phase encoding direction. The parameter *t* represents the scan time of the acquisition without oversampling or acceleration along kz for one phase encoding direction (c, i). R is the total under-sampling factor for each sampling pattern.

Moreover, due to the limited FOV and non-rectangular RF profile along the slice direction, where transition bands and side lobes extend to adjacent slabs (Fig. 2a), slice aliasing happens at slab boundaries in 3D multi-slab imaging. The slice aliasing artifacts can be reduced by over-sampling along kz with an extended FOV (Fig. 2b). To extend the FOV without increasing the number of shots (e.g., from 10 shots to 12 shots comparing Fig. 2c, i and Fig. 2c, ii), a higher acceleration factor along kz is required, which leads to a more challenging reconstruction.

Here, we combined blip-reversed acquisitions with two sampling strategies to improve the under-sampled reconstruction, which is particularly important for the first stage of reconstruction in which images are separately reconstructed for the blip-up and blip-down segments. First, kz blipped-CAIPI^25, 27^ was integrated for more effective use of coil sensitivity, where each shot covers the full extent of the phase encoding direction (ky) with even spacing along ky and includes “blips” to traverse closely spaced kz planes (Fig. 2c, iv). We also employed partial Fourier along kz (Fig. 2c, v), which reduces aliasing in the slice direction through a more densely sampled central k-space region. This achieves an even shorter scan time by reducing the number of shots (Fig. 2c, v, halved scan time compared to Fig. 2c, i). The proposed sampling approach enables simultaneous correction of distortions (by allowing joint acquisition of blip-up/down data) and boundary slice aliasing (by oversampling along kz) without increasing the scan time. We refer to our sampling approach as “CAIPI-PF” hereafter.

For the joint acquisition of blip-reversed data, equal numbers of shots traversing ky in opposite directions are acquired. The blip-up and blip-down sampling cover the complementary subsets of the k-space with a shift of Δ*k*_*y*_ and complementary partial Fourier along kz (Fig. 1, left), which reduces the noise amplification and prevents resolution loss in the stage 2 joint reconstruction.

#### 2.1.2. Reconstruction

The cost function of the SPIRiT-based regularized reconstruction is:

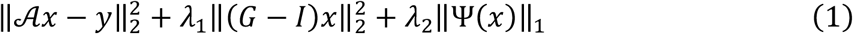

where x is the multi-coil k-space data of the target aliasing- and distortion-corrected image, 𝒜 is the forward operator, y is the acquired data, G is the SPIRiT kernel trained on coil calibration data, I is the identity matrix, Ψ is the wavelet operator, and *λ*_1_ and *λ*_2_ are the parameters for the SPIRiT and sparsity regularizations (the SPIRiT weights for stage 1 and stage 2 are denoted as *λ*_1,1_ and *λ*_1,2_, respectively). The SPIRiT regularization (*λ*_1_) facilitates parallel imaging by enforcing calibration consistency between every k space data point and its neighbors^26^, while the sparsity regularization (*λ*_2_) is used to suppress the noise^28^.

The forward operator 𝒜 is constructed differently for the two stages:

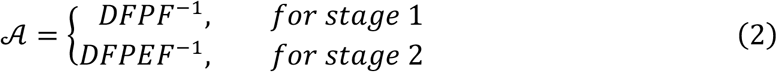

where D is the down-sampling operator in k-space, *F* and *F*^−1^ are the Fourier Transform and inverse Fourier Transform, respectively, and P represents motion-induced phase errors captured by 2D navigators. The E operator is only applied in stage 2 and represents spatial distortion induced by field inhomogeneity estimated from the stage 1 reconstruction.

It is worth noting that unlike the original SPIRiT formulation^26^ in which data consistency can be ensured implicitly by only estimating the missing k-space points, in our work the entire k-space needs to be estimated. This is because in our forward operator 𝒜 the acquired k-space points are corrupted by the motion-induced phase P and the distortion-induced displacement E, and are therefore different from those in the uncorrupted target k-space data x. Hence, the SPIRiT constraint is explicitly added as a regularization term and the data consistency term 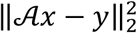 imposes the consistency constraint across all k-space points.

### 2.2. In vivo experiments

A 3D multi-slab spin-echo diffusion MRI sequence^8^ was modified to integrate blipped-CAIPI and kz partial Fourier sampling. After the imaging echo, a second refocusing pulse was applied to acquire a low resolution 2D navigator to correct the motion-induced phase errors. Subjects were scanned on a Siemens 7T scanner using a 32-channel receive coil. Written informed consent in accordance with local ethics was obtained from each subject.

#### 2.2.1. Evaluation protocols

To evaluate the impact of different sampling patterns and reconstruction parameters, fully sampled single-slab datasets using CAIPI sampling were acquired from a single subject using the following scan parameters: 1.22mm isotropic resolution, 20 slices, TE1(imaging)/TE2(navigator)/TR=72/128/1800ms, b=1000s/mm^2^, diffusion encoding along left-right direction, ky under-sampling Ry=3, echo spacing 0.76ms, navigator size: 64 phase encoding lines. To avoid slice aliasing, 1.2x oversampling along kz was applied, which encoded a larger FOV of 29.3mm along the slice direction with 24 slices. Three scans with 0, 1, and 2Δ*k*_*y*_ shift were acquired and combined to produce fully sampled data. Blip-up and blip-down phase encoding datasets were acquired separately. In addition, a dataset using conventional rectangular sampling without kz oversampling was also acquired from the same subject to demonstrate slice aliasing.

Retrospective under-sampling was then performed to evaluate different sampling patterns, which were similar to those shown in Fig. 2 but with 20 or 24 kz encoding planes (without or with kz oversampling). For CAIPI-PF sampling (Fig. 2c, v), the shots #1,3,5,7,9,11,12,13,14,15 were used for acquiring blip-up data and the shots # 11,12,13,14,15,16,18,20,22,24 were used for acquiring blip-down data (shot #n covers kz plane n and n+1) (partial Fourier factor: *f*_*pf*_ = 2/3).

#### 2.2.2. High-resolution diffusion protocols

Six subjects were scanned using a 1.05mm isotropic resolution protocol to demonstrate the robustness of the proposed method. Six slabs with 20 slices per slab were acquired. Neighboring slabs were overlapped by 1 slice, resulting in 115 slices in the final reconstruction. The FOV was 220×220×121 mm^3^ and voxel size was 1.05mm isotropic. Interleaved slab acquisition was used to minimize cross talk between adjacent slabs. Diffusion-weighted images were acquired with b=1000s/mm^2^ and 48 diffusion directions uniformly sampled on a sphere with interleaved 3 b=0 image volumes. The echo spacing was 0.82 ms, and Ry=3 acceleration was applied along ky phase encoding, resulting in an effective echo spacing of 0.27 ms. TE1(imaging)/TE2(navigator)/TR=82/150/1800ms. The diffusion weighted data were acquired using the same CAIPI-PF sampling as in the evaluation protocol (i.e., Fig. 2c, v) (*f*_*pf*_ = 2/3) with 10 blip-up shots and 10 blip-down shots for each slab. To reduce aliasing artifacts from CSF signal, b=0 images were acquired with kz fully sampled (i.e., 24 blip-up shots and 24 blip-down shots for each slab). The total scan time was ∼33min (∼36s per diffusion direction).

In one subject, a dataset using a conventional 3D multi-slab high-resolution protocol^8^ was also acquired using 20 kz rectangular sampling (Fig. 2c, i) to enable direct comparison. Eight b=0 images, including 6 blip-up and 2 blip-down volumes, were interspersed into the diffusion-weighted image acquisition. Other scan parameters were the same as the CAIPI-PF protocol and the total scan time was ∼34min.

An MPRAGE image was also acquired for each subject as an anatomical reference (∼5 min) at 0.86mm isotropic resolution.

### 2.3. Reconstruction details

The integrated blip-reversed 3D multi-slab EPI data were reconstructed using the proposed two-stage reconstruction, while the conventional 3D multi-slab EPI data were reconstructed by the stage 1 reconstruction. The SPIRiT kernel used in the reconstruction was trained using gradient echo coil calibration data. The image reconstruction was conducted offline in MATLAB. Image processing were conducted using functions from FMRIB Software Library (FSL)^23^ unless indicated otherwise. Eq. 1 was optimized with preconditioned conjugate gradient method with variable splitting^29^. The k-space data were first Fourier transformed along kx followed by reconstruction performed for each *k*_*y*_-*k*_*z*_ plane using a 5×5 SPIRiT kernel. The reconstructed 2D images were concatenated along the readout direction (x) to form the whole image volume. The 32-channel data were compressed to 8 to shorten the calculation time^30^. The field map was calculated using “topup” ^13, 24^. On a 2.9 GHz Quad-Core Intel Core i7 CPU, the computation time for one *k*_*y*_-*k*_*z*_ plane of one slab is ∼15 seconds for stage 1 reconstruction, and ∼90 seconds for stage 2 reconstruction. The processing time for “topup” is ∼5 minutes per slab.

In stage 1 reconstruction, the unacquired partial Fourier region was zero filled before inverse Fourier transform along kz, resulting in smoothing along the slice direction. As the ΔB0 field is spatially smooth and the slab is thin, the impact of slice smoothness on field map estimation is expected to be not significant. No zero filling is necessary for the stage 2 reconstruction because it operates on the full kz extent (complementary blip-up and blip-down segments).

The 2D navigator images were reconstructed with 2D GRAPPA^31^ using a 2D calibration dataset, which was acquired separately for blip-up and blip-down data^8^. The phase images were extracted and used as an estimation of motion induced phase errors.

To find the optimal SPIRiT regularization parameter *λ*_1_, reconstruction of CAIPI-PF sampled data was performed with different *λ*_1_ values. Specifically, *λ*_1,1_ = 0,0.1,1,10 and *λ*_1,2_ = 0,1,20,100 were assessed for stage 1 and stage 2, respectively, using the data acquired with the evaluation protocol. The optimal *λ*_1,1_ for stage 1 reconstruction was used to calculate blip-up/down images and estimate a field map, which were then used for *λ*_1,2_ optimization of stage 2 reconstruction. The similarity between the reconstructed image with the fully sampled reference was evaluated using normalized root mean squared error (NRMSE). The optimal SPIRiT regularization parameters, *λ*_1,1_ = 1 for stage 1 and *λ*_1,2_ = 20 for stage 2, were used for the reconstruction of high-resolution datasets. The sparsity regularization parameter *λ*_2_ = 0.7 × 10^−1^ was used in both stage 1 and stage 2 reconstructions. The estimated field maps were compared with reference field maps derived from fully sampled blip-up and blip-down images, and voxel displacement errors were calculated by multiplying the field map difference (in Hz) with the readout duration (in seconds). NRMSE and mean voxel displacement errors were calculated within a brain mask.

The g-factors for different blip-up sampling patterns demonstrated in Fig. 2 were calculated using the pseudo-multiple replica method^32^ with 100 repetitions of the Monte-Carlo simulation. In each Monte-Carlo repetition, independent and identically distributed complex Gaussian noise was added to the pre-whitened multi-channel k-space data. For the stage 1 CAIPI-PF sampling, the impact of kz partial Fourier on g-factor was accounted by dividing the resultant g-factor with the square root of partial Fourier factor as suggested by Kettinger et al^33^. The sparsity constraint was not included in g-factor calculations to reduce non-linearity. The g-factors were calculated within a brain mask.

### 2.4. Post-processing

Slab combination and correction for slab saturation artifacts were performed using “NPEN”^20^. The whole-brain images were corrected for Gibbs ringing using “mrdegibbs3D” (https://github.com/jdtournier/mrdegibbs3D)^34, 35^. For conventional 3D multi-slab data acquired with the high-resolution protocol, a whole-brain field map was estimated using blip-up and blip-down b=0 image volumes using “topup”, which was then input to FSL’s “eddy”^36^ along with all diffusion data to correct for susceptibility and eddy current induced distortions. The CAIPI-PF data acquired with the high-resolution protocol were processed with “eddy” without “-topup” option, as field inhomogeneity-induced distortion has been corrected in the reconstruction. To evaluate the efficacy of the proposed method in estimating diffusion direction-dependent dynamic field maps and reducing the eddy current induced geometric distortions, the CAIPI-PF data and conventional 3D multi-slab data were also processed by “eddy” without eddy current correction, which was facilitated by specifying “— flm=movement” in the command line of “eddy”.

### 2.5 Diffusion analysis

All diffusion analyses were performed in the native diffusion space. For the subject scanned twice using the CAIPI-PF and conventional 3D multi-slab high resolution protocol, results were co-registered to the anatomical space for comparison. These datasets were acquired in separate sessions due to the long scan times. The diffusion data from both samplings were co-registered to the MPRAGE image. Due to different head positions, distortions induced by gradient nonlinearity were different between the two scans. For fair comparison, the MPRAGE image was acquired in a separate session along with an intermediate b=0 image volume with matched gradient nonlinearity distortions to minimize the impact of gradient nonlinearity and improve the co-registration accuracy. These co-registration steps are listed in the Co-registering Details section in Supplementary Information.

For the remaining subjects, T1w images were co-registered to the diffusion space. The b=0 images were co-registered to the T1w images using “epi_reg”^37, 38^. The resultant transformations were then inverted using “convert_xfm” ^37, 38^ and applied to the T1w images.

The diffusion tensor model fitting was performed using “dtifit”^24^. White matter tractography was performed using “autoPtx”^39-41^, which includes the pre-processing stage that runs probabilistic model fit using “bedpostx”^40^, and the tractography stage which runs the probabilistic tractography using “probtrackx”^40^.

## 3. Results

Figure 3 shows the effectiveness of the proposed sampling in reducing the slice aliasing in the slab boundary slice. Without oversampling, the boundary slice suffers from aliasing (comparing Fig. 3, i with Fig. 3, iv) caused by non-rectangular RF profile and limited FOV (Fig. 2a). Simple oversampling along kz increases the FOV (Fig. 2b) and corrects the aliasing (comparing Fig. 3, ii with Fig. 3, iv), but requires longer scan time. Our proposed CAIPI-PF sampling produces boundary slice aliasing-corrected images with a much faster acquisition, reducing scan time from 1.2t to 0.5t (t represents the scan time of a rectangular sampling without over-sampling or acceleration along kz, as Fig. 2c, i).

**Figure 3.**
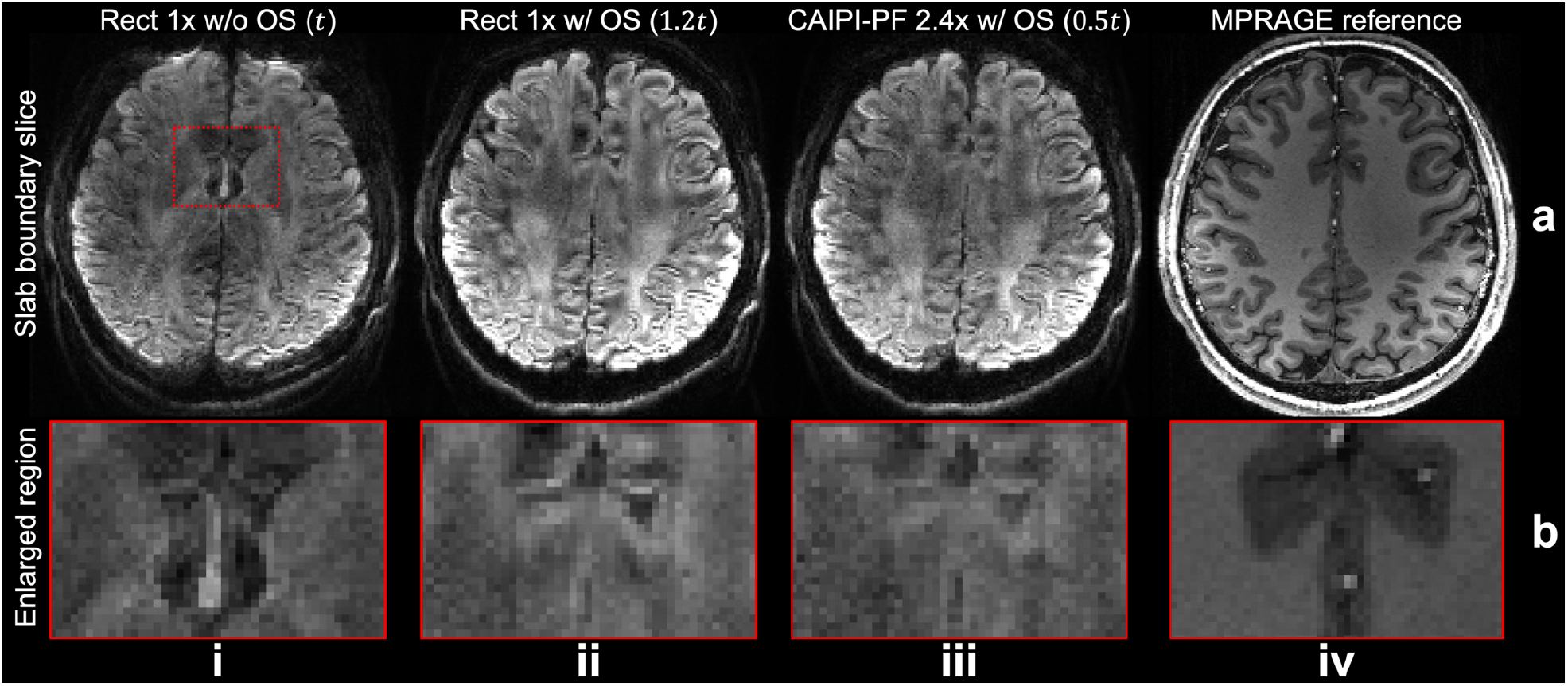
Boundary slice aliasing correction. The boundary slices (a) and enlarged regions (b) of blip-up data from rectangular sampling without oversampling along kz (Rz/Ry=1/3) (i), rectangular sampling with 20% oversampling and 2x acceleration along kz (Rz/Ry=1/3) (ii), the proposed CAIPI-PF sampling with 20% oversampling, partial Fourier and 2.4x acceleration along kz (Rz/Ry=2.4/3) (iii) acquired with the evaluation protocol (1.22 mm isotropic resolution), and the reference MPRAGE data (0.86 mm isotropic resolution) (iv) are displayed, with relative scan times listed for each method except for MPRAGE reference. The parameter *t* represents the scan time of a rectangular sampling without over-sampling or acceleration along kz (as Fig. 2c, i).

Results from stage 1 blip-up reconstruction for different sampling patterns are demonstrated in Fig. 4. The reference data were fully sampled with no acceleration along ky or kz. The results with fully kz encoding and Ry=3 under-sampling along ky produces the most similar result to the reference and the lowest g-factor (NRMSE: 9.3%; g-factor: 1.12) with the longest acquisition time. Image from rectangular under-sampling with Ry/Rz=3/2 suffers from strong aliasing due to the limited coil sensitivity variation along the thin slab, producing the highest error and g-factor (NRMSE=27.1%; g-factor: 1.75). Using CAIPI sampling alone can substantially improve the reconstruction and lower the NRMSE (16.4%) and g-factor (1.45) by more efficient use of coil sensitivity information. The partial Fourier strategy further reduces reconstruction errors with lower NRMSE (15.6%) and g-factor (1.24) using an even shorter scan time. The blip-down reconstruction produces consistent results with the blip-up reconstruction (Fig. S1).

**Figure 4.**
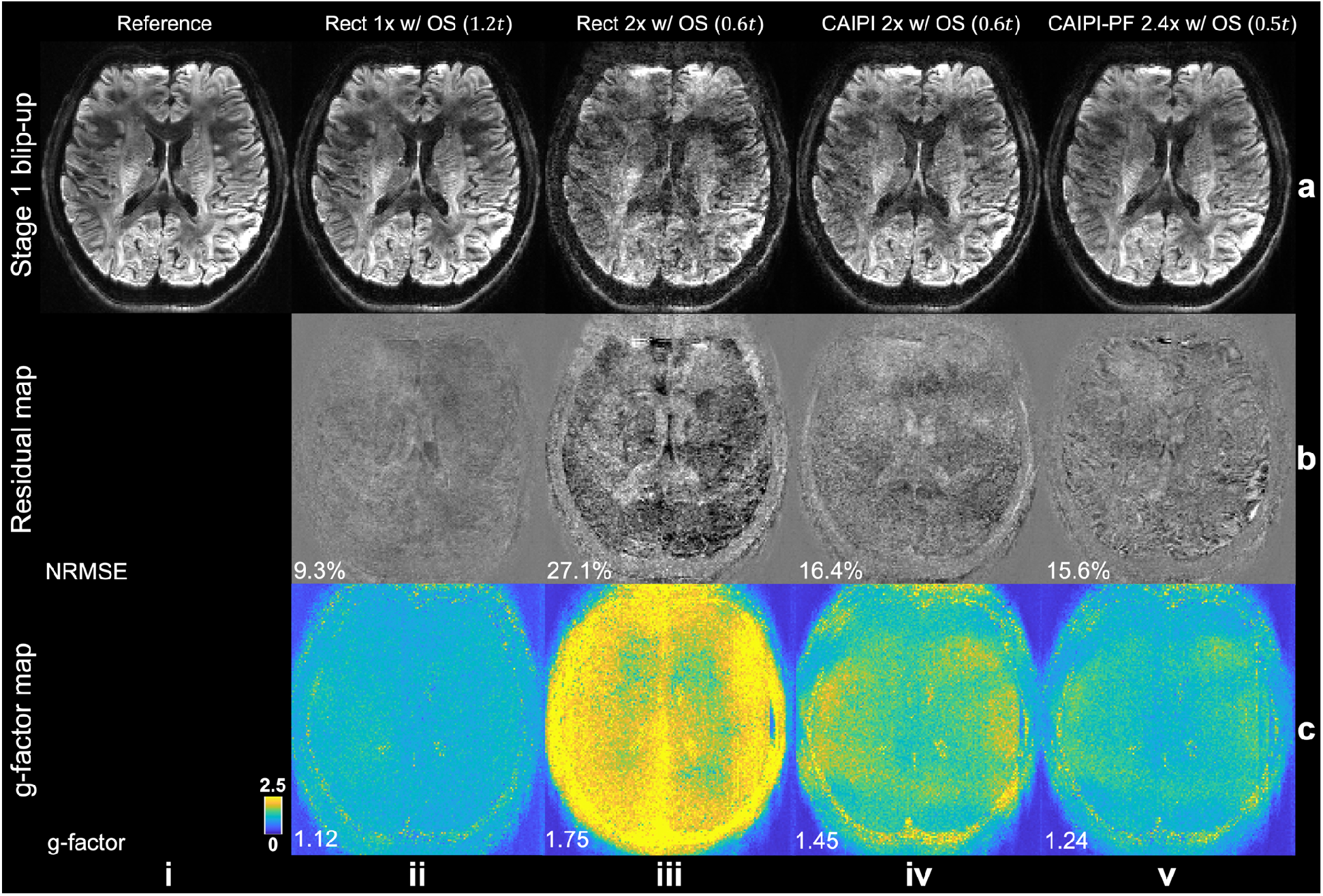
Stage 1 blip-up reconstruction. The blip-up data for a slice at slab center (a) from fully sampled reference (i) and different under-sampling patterns: rectangular sampling with 20% oversampling along kz (Rect 1x w/ OS) (ii); rectangular sampling with 20% oversampling and 2x acceleration along kz (Rect 2x w/ OS) (iii); CAIPI sampling with 20% oversampling and 2x acceleration along kz (CAIPI 2x w/ OS) (iv); CAIPI-PF sampling with 20% oversampling, partial Fourier and 2.4x acceleration along kz (CAIPI-PF 2.4x w/ OS) (iv), their residuals with the fully sampled reference (b), and their g-factor maps (c) are displayed, with relative scan times listed for each method. The data were acquired with the evaluation protocol (1.22 mm isotropic resolution). The normalized root mean squared errors (NRMSE) and mean g-factor of the whole slab are listed to quantify the image similarity and noise amplification. The parameter *t* represents the scan time of a rectangular sampling without over-sampling or acceleration along kz (as Fig. 2c, i).

The estimated field maps and stage 2 joint reconstruction results for different joint blip-reversed acquisition strategies are shown in Fig. 5. The data acquired with full kz sampling still produces the most accurate field map and the best stage 2 reconstruction (mean displacement error: 0.17 pixels; NRMSE: 6.6%). However, the scan time of this acquisition strategy is 2.4x the scan time of a conventional multi-slab acquisition. In conventional rectangular under-sampling, the reconstructed image exhibits obvious structural difference with the reference due to the inaccurate field map estimated from the aliased images from stage 1 reconstruction (mean displacement error: 1.09 pixels; NRMSE: 30.2%). The residual aliasing of regular CAIPI under-sampling in stage 1 reconstruction results in errors in the estimated field map, which leads to inaccuracies in joint reconstruction results (mean displacement error: 0.35 pixels; NRMSE: 14.2%). The proposed CAIPI-PF sampling provides an improved field map estimation and joint reconstruction with substantially better results due to a more densely sampled k-space center (mean displacement error: 0.24 pixels; NRMSE: 12.6%) using the shortest scan time among all sampling patterns. The voxel displacement error map from CAIPI-PF also shows similar spatial pattern to the fully kz-sampled data without obvious anatomical bias, indicating the CAIPI-PF sampling strategy does not introduce substantial errors in field map estimation.

**Figure 5.**
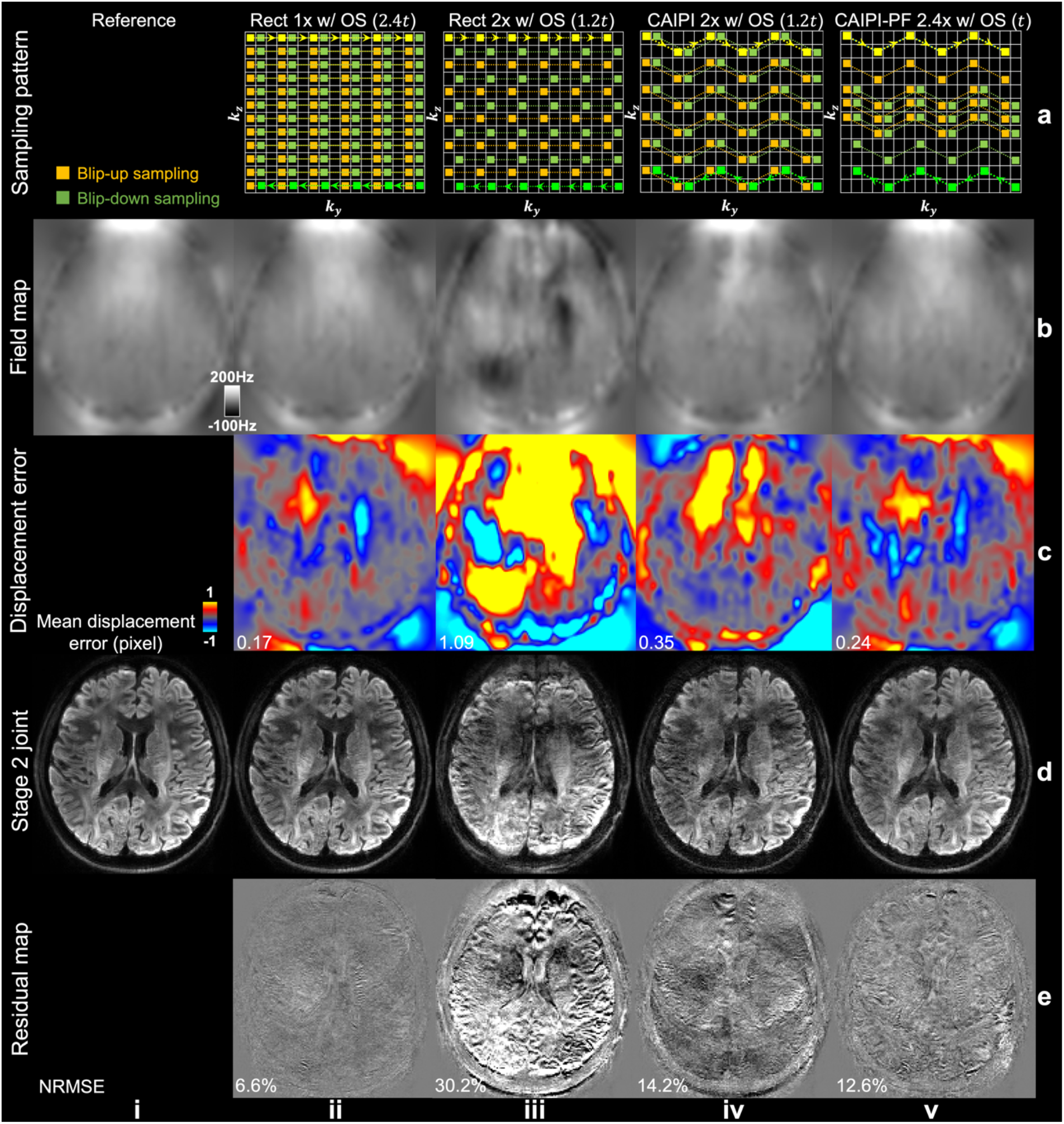
Field map estimation and joint reconstruction. The joint blip-up (yellow) and blip-down (green) sampling patterns (a), the corresponding estimated field maps from “topup” (b), the voxel displacement error maps (c), stage 2 joint reconstruction results (d), and residual maps (e) with the reference image acquired with the evaluation protocol (1.22 mm isotropic resolution) are displayed, with relative total scan times listed for each method. The parameter *t* represents the scan time of a rectangular sampling without over-sampling or acceleration along kz for one phase encoding direction (as Fig. 2c, i). The trajectory of one shot of blip-up and blip-down sampling is marked in bright yellow and bright green, respectively. The mean displacement errors and normalized root mean squared errors (NRMSE) of the whole slab are listed to quantify the image similarity.

Figure 6 characterizes the impact of SPIRiT regularization parameter *λ*_1_ on the reconstruction of CAIPI-PF data for both stages of reconstruction. Because of the different sampling patterns and forward models for stage 1 and stage 2 reconstruction, *λ*_1_ was separately optimized for two stages (*λ*_1,1_ = 0,0.1,1,10 and *λ*_1,2_ = 0,1,20,100 were evaluated for stage 1 and stage 2, respectively). Without SPIRiT constraint (i.e., *λ*_1_ = 0), the reconstruction fails to leverage the redundant coil information and is extremely ill-posed, leading to an aliased reconstruction with the highest NRMSE (40.5% for stage 1, 33.5% for stage 2). SPIRiT constraints substantially reduce the aliasing and improve the reconstruction. The increase of SPIRiT weights leads to a higher-SNR but slightly more biased output images. Therefore, moderate SPIRiT weights *λ*_1,1_ = 1 for stage 1 and *λ*_1,2_ = 20 for stage 2 are selected for our current reconstruction, which produces the lowest NRMSE (15.6% for stage 1 and 12.6% for stage 2). Their residual maps do not contain obvious anatomical difference, suggesting no significant blurring or bias effect. The impact of the stage 1 SPIRiT weight on the field map estimation is consistent with its impact on the blip-up image quality (Fig. S2).

**Figure 6.**
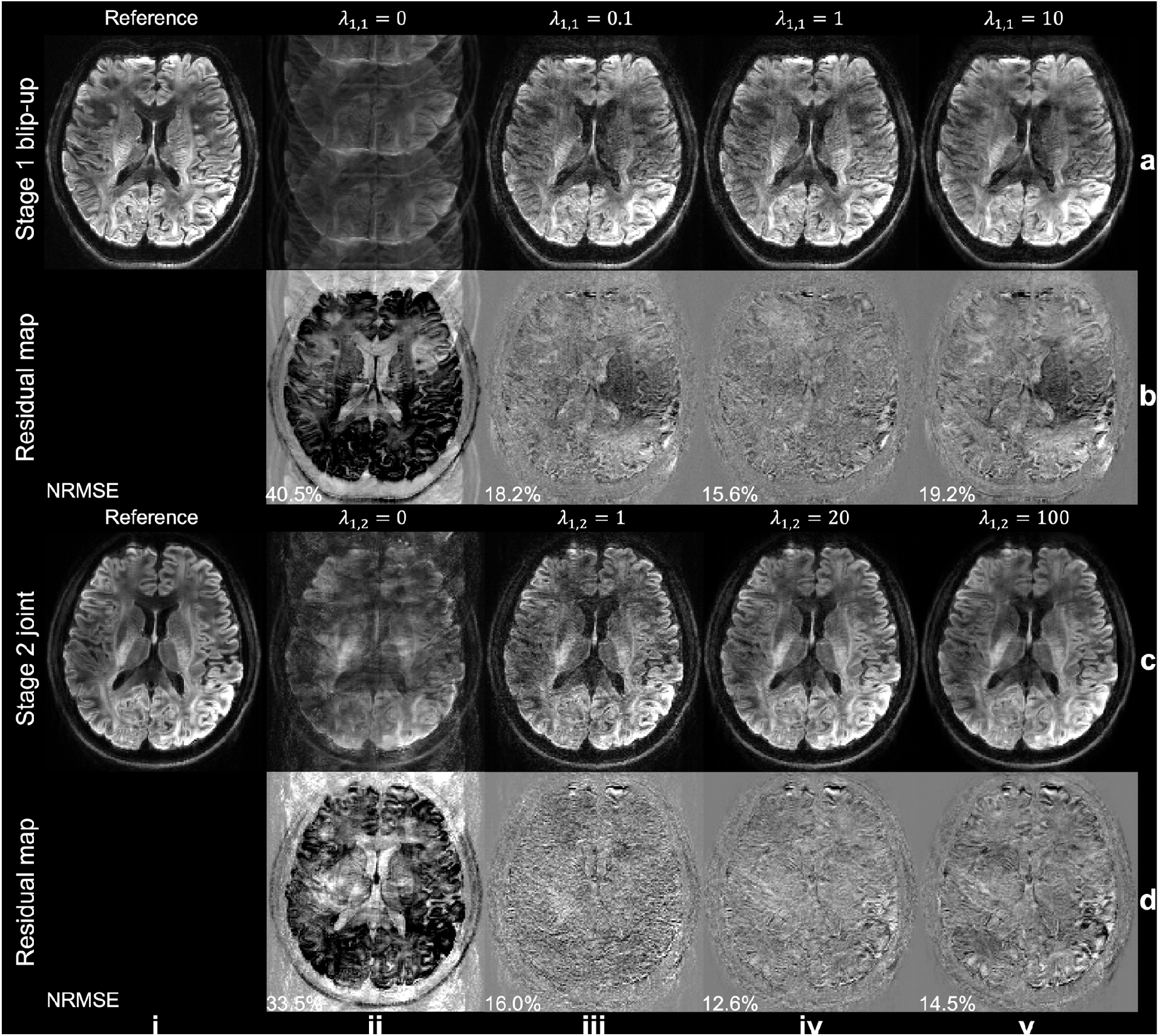
Impact of SPIRiT regularization weight on reconstruction. Stage 1 blip-up (a) and stage 2 joint reconstruction (c) of CAIPI-PF data using different SPIRiT regularization weights and their residuals compared to the reference (b, d) acquired with the evaluation protocol (1.22 mm isotropic resolution) are displayed. The optimal weight for stage 1 (i.e., *λ*_1,1_ = 1) was used when different weights were evaluated for stage 2. The normalized root mean squared errors (NRMSE) of the whole slab are listed to quantify the image similarity.

Results from the 1.05 mm whole-brain high-resolution protocols are showed in Fig. 7. Our proposed method produces single-volume diffusion-weighted image with high SNR and reduced artifacts and high-quality whole-brain diffusion tensor imaging (DTI) results. Compared to the conventional 3D multi-slab acquisition where diffusion weighted images were acquired with only blip-up phase encoding, and distortion correction was applied using a field map derived from a pair of blip-up/down b=0 image, the proposed method achieves substantially improved anatomical fidelity, especially near the regions in the frontal lobe and pons where the field inhomogeneity is strong (Fig. 7a, yellow arrows). The conventional method also suffers from residual slice aliasing at slab boundaries (Fig. 7c, ii, the white arrow in the splenium of corpus callosum) due to the absence of over-sampling and limited overlapping between slabs (only 1 slice overlapping) to match the scan time with the proposed method.

**Figure 7.**
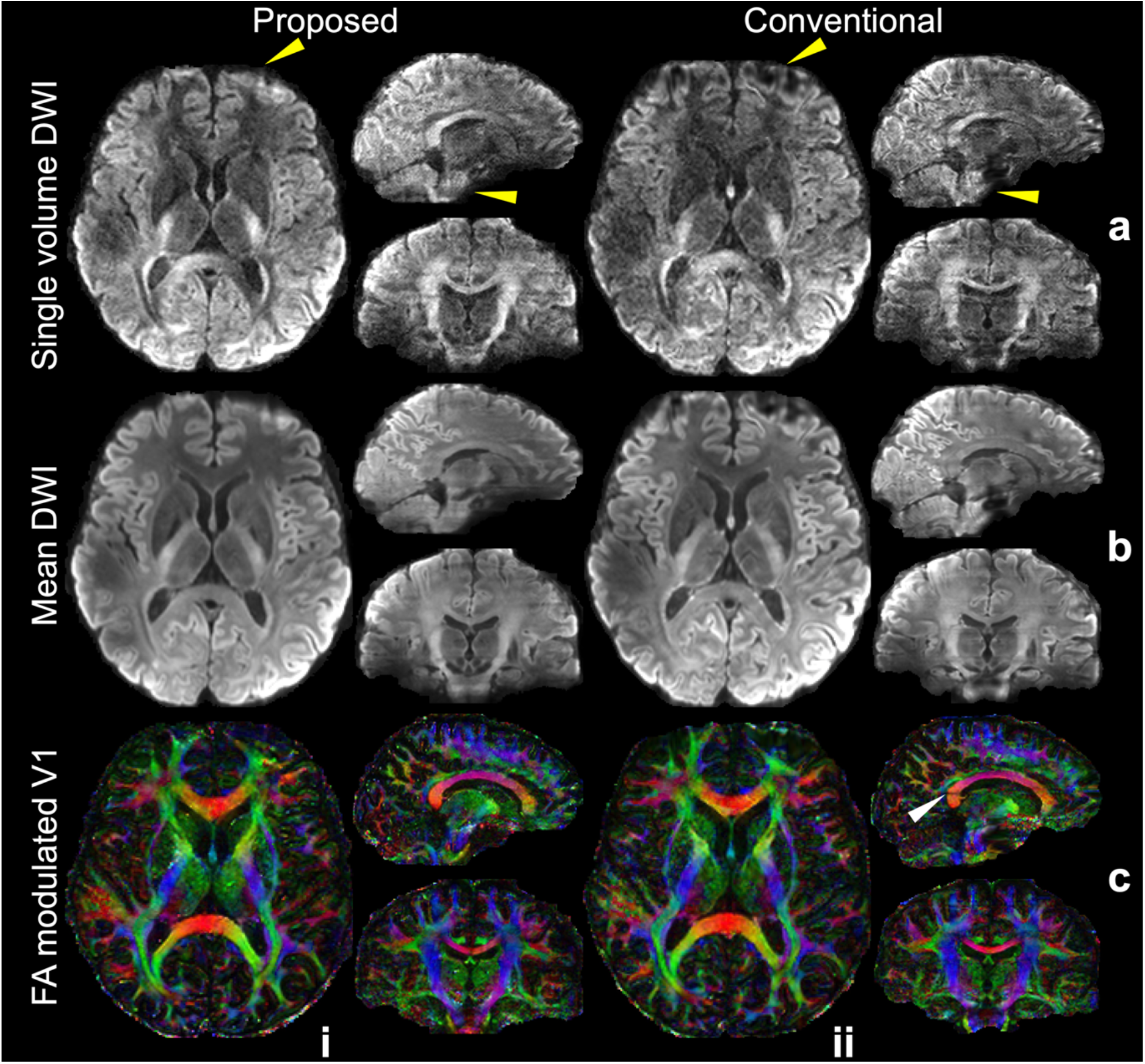
High resolution whole-brain diffusion images. The single volume diffusion-weighted images (DWI) along the diffusion direction (−0.10, 0.88, 0.47) (a), the mean DWI (b), and the fractional anisotropy (FA) modulated primary eigenvector (V1) (c) of diffusion data acquired using joint blip-reversed with CAIPI-PF sampling (Proposed, i) and conventional 3D multi-slab sampling using only blip-up phase encoding (Conventional, ii) at 1.05 mm isotropic resolution from the same subject are displayed. The diffusion data are co-registered to the same anatomical data (0.86 mm isotropic resolution) for comparison. The yellow arrows highlight the regions with strong distortions near frontal and pons (a), and the white arrow highlights the boundary slice aliasing (c), which cannot be fully corrected with the conventional method.

The alignment of the diffusion data with the anatomical reference is showed in Fig. S3. When combining blip-up and blip-down data for distortion correction, the proposed method and the conventional method produce similar distortion correction on b=0 images. However, the distortion-corrected diffusion-weighted images from the proposed method exhibits consistently improved anatomical fidelity for different diffusion directions. In addition, the proposed method can also correct eddy current induced distortions because the diffusion direction-dependent dynamic field map is incorporated into the joint reconstruction (Fig. S4). The co-registered images with and without eddy current correction are highly similar from the proposed reconstruction, with a low mean NRMSE across all diffusion directions (1.7%), indicating most eddy current induced distortions have already been corrected during the reconstruction. In comparison, images from the conventional method exhibit larger differences before and after the eddy current correction, with a much higher mean NRMSE for all diffusion directions (9.4%).

Tractography results show consistent improvement of the proposed method compared to the conventional method. Improvements can be seen in frontal regions with residual distortion for the anterior thalamic radiation (Fig. 8a) and forceps minor (Fig. 8b), where the proposed method captures projections into the cortical gray matter more successfully. Similarly, the corticospinal tracts are represented as thicker white matter bundles in tractography based on the proposed reconstruction compared to the conventional method, whose thickness is considerably reduced near the pons (red arrows, Fig. 8c). The tract mask (binarized with tract density threshold: 0.3%) volumes from the proposed method increase by 20.2%, 43.0%, and 49.5% compared to those from the conventional method for right anterior thalamic radiation, forces minor, and right corticospinal tracts, respectively. The detailed tract mask volumes can be found in Table S1.

**Figure 8.**
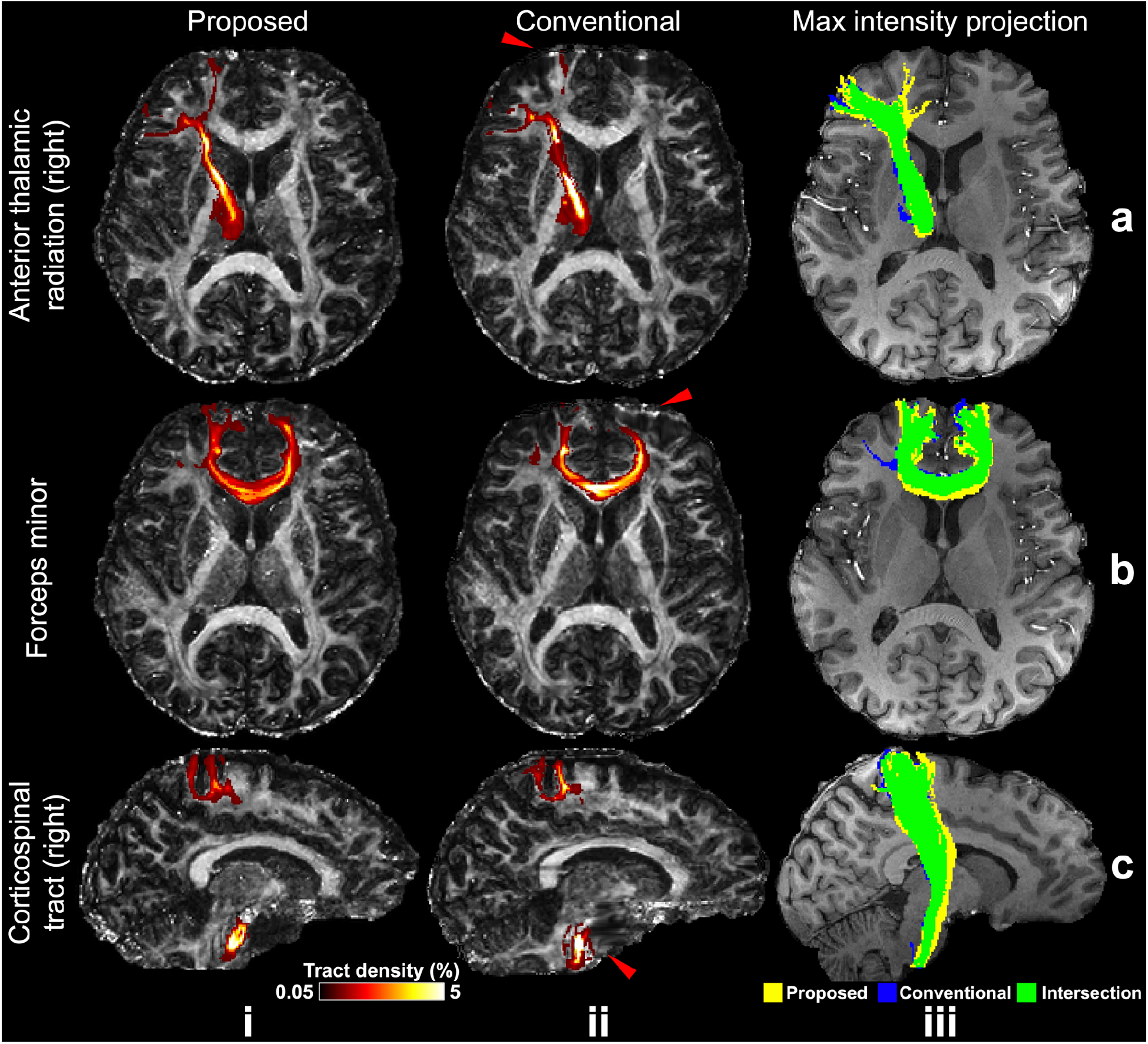
Comparison of tractography results. Tractography results of right anterior thalamic radiation (a), forces minor (b), right corticospinal tract (c) from the same subject using the proposed joint blip-reversed with CAIPI-PF (Proposed, i) and conventional 3D multi-slab sampling (Conventional, ii) with high-resolution protocols (1.05 mm isotropic) overlayed on their fractional anisotropy (FA) maps (tract density displaying range: 0.05%-5%), and their maximum intensity projection masks (iii) (binarized with tract density threshold: 0.3%) overlayed on the anatomical image. The red arrows highlight the regions with strong distortions near frontal (a, b, ii) and pons (c, ii), which cannot be fully corrected with the conventional method.

These results generalize to other subjects (Fig. 9). The single volume diffusion-weighted images show high-SNR and sharp textures. The DTI maps demonstrate high-quality and resolve fine structures with the high-resolution. The mean DWI results demonstrate high anatomical fidelity, with high structural similarity compared to the anatomical reference. Results of two other subjects are shown in Fig. S5.

**Figure 9.**
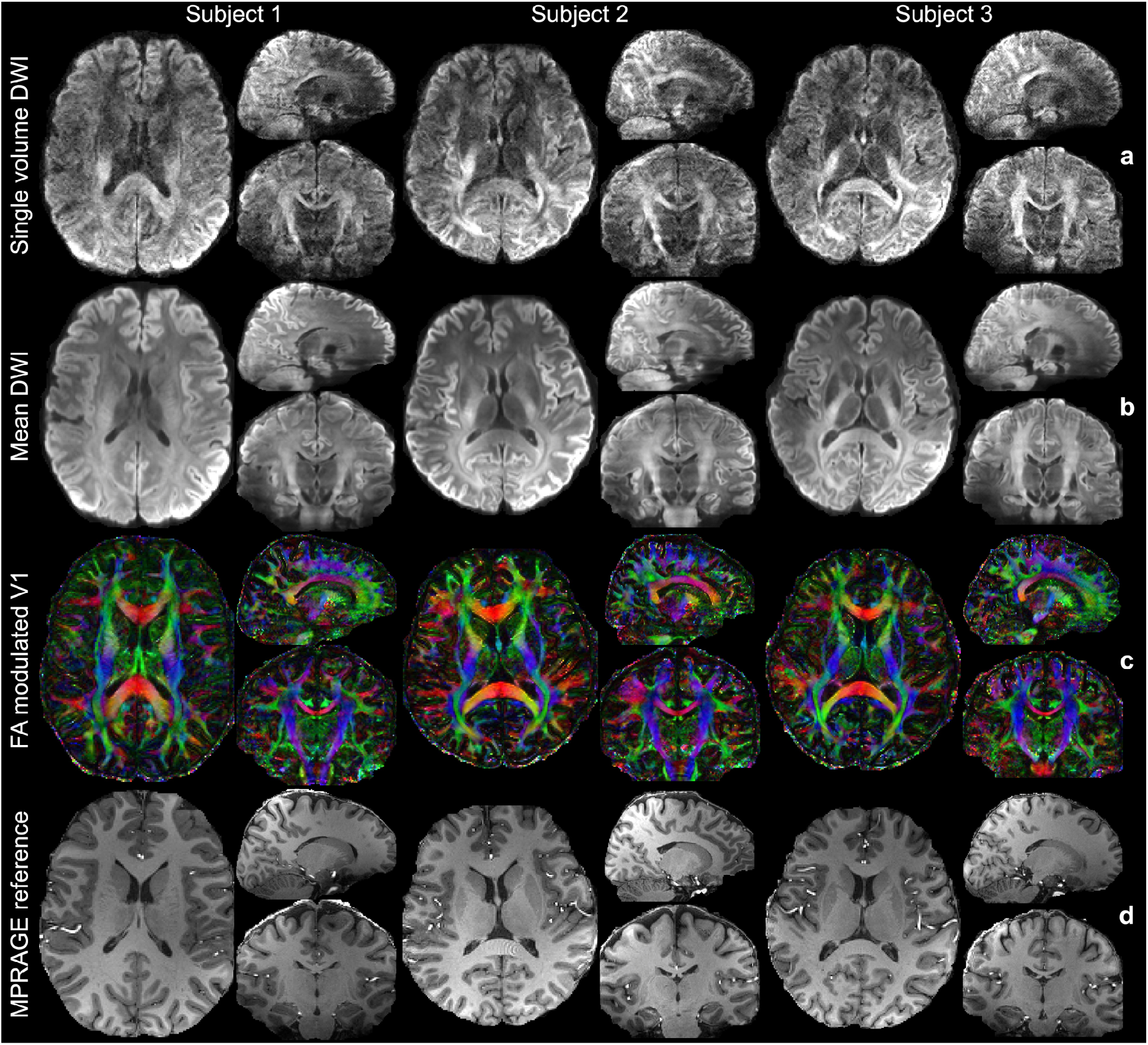
Diffusion MRI results of multiple subjects using the proposed method. The single volume diffusion-weighted images (DWI) along the diffusion direction (−0.26, −0.81, −0.52) (a), the mean DWI (b), and the fractional anisotropy (FA) modulated primary eigenvector (V1) of diffusion tensor (c), and the anatomical images for reference (d) of three subjects are displayed. The diffusion images are acquired with the high-resolution protocol at 1.05 mm isotropic resolution using joint blip-up/down with CAIPI-PF acquisition. The anatomical images are acquired with MPRAGE at 0.86mm isotropic resolution and co-registered to diffusion space for comparison.

Tractography results for these subjects are shown in Fig. 10. The anterior thalamic radiation and forceps minor project successfully into the cortical gray matter in regions of strong field inhomogeneity. The corticospinal tract is well reconstructed even in inferior regions such as the pons where B0 field inhomogeneity is strong. The tractography results for these slices without maximum intensity projections are shown in Fig. S7.

**Figure 10.**
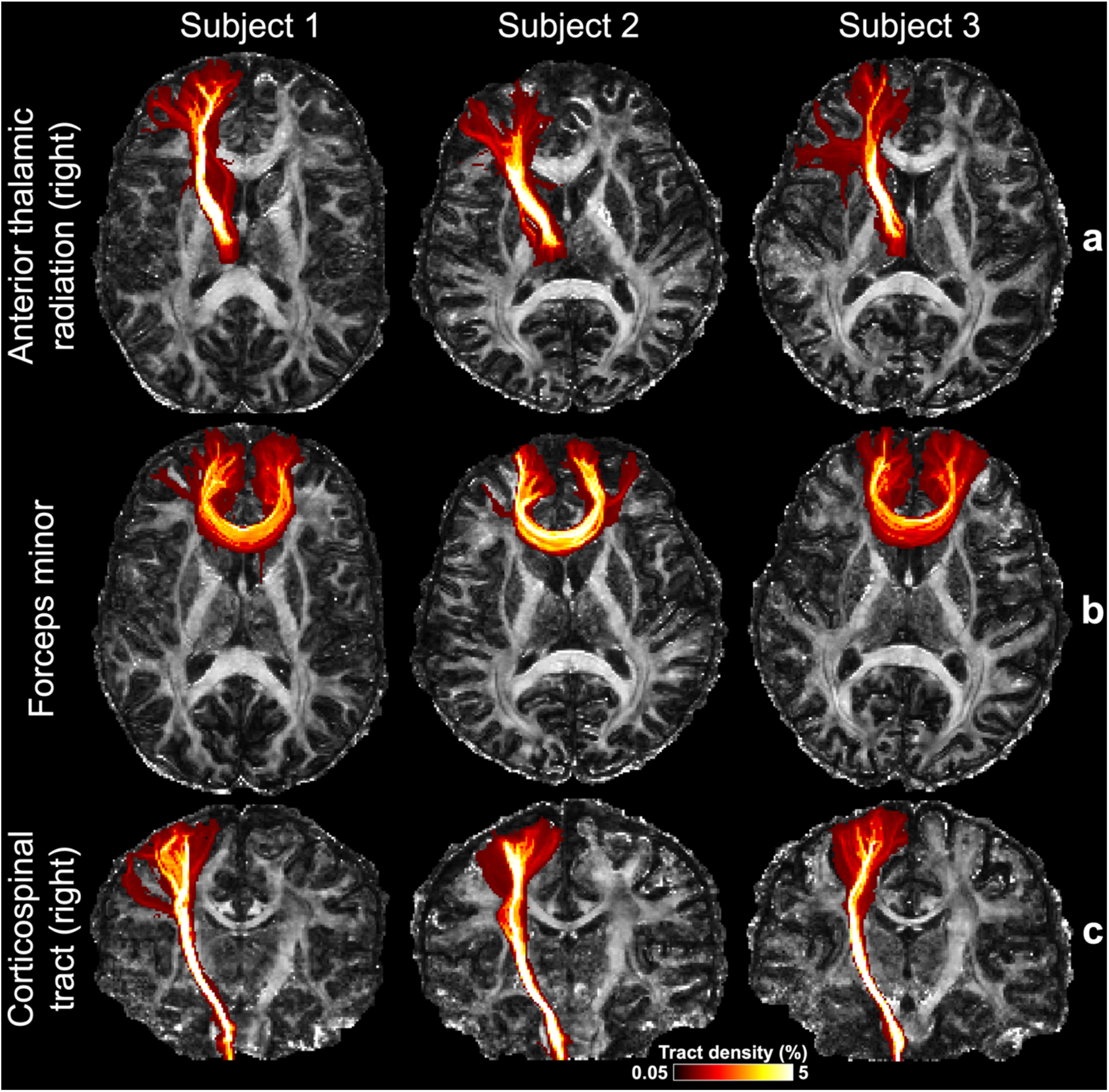
Maximum intensity projections of fiber tracking results from three subjects. Three tracts including right anterior thalamic radiation (a), forceps minor (b), and right corticospinal tract (c) derived from data acquired with the high-resolution protocol from three subjects using the joint blip-reversed CAIPI-PF acquisition are displayed, overlayed on their fractional anisotropy (FA) maps. All tracts are visualized using the same track density threshold (0.05%-5%).

## 4. Discussion

In this work, we demonstrate that high resolution 3D multi-slab diffusion MRI with minimal distortion and slice aliasing can be achieved by incorporating multiple sampling strategies into a joint reconstruction framework. For accurate distortion correction, both blip-up and blip-down shots for each diffusion direction were acquired for field mapping and distortion suppression. Over-sampling along kz was used for an extended FOV to minimize slice aliasing at slab boundaries. Blip-reversed based distortion correction and boundary slice aliasing correction have been proposed previously^9, 17-20^. One of the key challenges in this work is to acquire blip-reversed data with kz over-sampling without requiring longer scan time than the conventional 3D multi-slab diffusion acquisition. We addressed this challenge using two alterations in the sampling pattern. First, blipped-CAIPI makes more efficient use of coil sensitivity and improves the conditioning of reconstruction. This is essential for 3D multi-slab imaging as each slab is very thin (∼20 mm) and variation of coil sensitivity across the slab is extremely limited. Without CAIPI the reconstruction is highly ill-posed and lead to high noise amplifications (Fig. 4, iii). Blipped-CAIPI uses the in-plane coil profiles to encode information, improving the conditioning of the reconstruction and reducing the g-factor significantly. Second, our kz partial Fourier strategy enables a denser sampling of the k-space center compared to regular under-sampling, which reduces the aliasing artifacts. The improvement is less obvious in stage 1 reconstruction (Fig. 4, iv, v), but is substantial in terms of the accuracy of the estimated field map and the image quality of stage 2 joint reconstruction (Fig. 5, iv, v). The resultant blurring along the slice direction in stage 1 reconstruction (residual structures in Fig. 4b, v) introduced by zero-filling of the partial Fourier region does not have an obvious impact on the accuracy of the estimated field map, presumably because field maps are intrinsically smooth, especially along the slice direction for thin slabs. In the joint reconstruction, such blurring is reduced thanks to the complementary patterns of blip-up and blip-down sampling.

The SPIRiT constraint used in this work improves SNR while preserving the sharp textures (Fig.6) and does not induce obvious blurring effect even when the regularization weights are relatively high (Fig.6, v). However, in this case the data consistency term would be compromised, leading to increased reconstruction errors. Therefore, moderate SPIRiT weights (i.e., *λ*_1,1_ = 1 for stage 1 and *λ*_1,2_ = 20 for stage 2) were used in our reconstruction. The low g-factors of our reconstruction probably result from the implicit conditioning of noise for k-space-based reconstruction^32^ and the noise averaging effect of SPIRiT regularization^26^. Additionally, our reconstruction can be highly parallelized because we used 2D SPIRiT (i.e., separately reconstructing each *k*_*y*_ − *k*_*z*_ plane) in our framework. Using 3D SPIRiT may potentially further improve the reconstruction by leveraging data redundancy along the readout direction, but at a cost of longer computation time. Future work will be needed to explore the performance of 3D SPIRiT on the 3D multi-slab data.

It is worth emphasizing the importance of correcting the slab boundary slice aliasing in our framework. Our stage 2 joint reconstruction relies on an accurate field map to produce distortion-corrected reconstruction for each slab, which will be affected by boundary slice aliasing in stage 1 reconstruction. Therefore, removing boundary slice aliasing with over-sampling along kz is essential for both aliasing-free reconstruction and accurate field map estimation.

The distortion correction efficacy of the proposed method is largely determined by the accuracy of field map estimation from stage 1 reconstruction. It is helpful to consider the “topup” correction using fully sampled blip-up and blip-down data as an upper bound for performance. In extreme cases when distortion is too strong for “topup” to correct even with fully sampled blip-up and blip-down data, the proposed method would have limited performance similar to “topup” correction. This is reflected in the results of one subject (Fig. S6, Subject 4), where residual distortions in the frontal region remain even after the joint reconstruction, probably due to the extremely high initial distortion level. Nevertheless, the level of residual distortion in the proposed method is highly similar to the “topup” corrected b=0 image with fully kz sampling (Fig. S6), indicating that the main source of the residual distortions is the extreme field inhomogeneity, which was likely due to poor shimming for that particular scan. A potential method to improve this is to use additional hardware to provide more homogeneous B0 field (e.g., using local shimming coils^18^).

## 5. Conclusion

In this work, an acquisition and reconstruction framework are developed to minimize the distortion and boundary slice aliasing in high-resolution 3D multi-slab diffusion MRI without increasing the scan time. The designed method achieves high-fidelity, robust reconstruction and produces high-SNR, high-quality diffusion images with superior anatomical fidelity compared to the conventional 3D multi-slab acquisition approach. The value of the method is demonstrated in improving the DTI fitting and tractography and can be further explored in more types of applications and by pushing to higher, submillimeter isotropic resolutions.

## Acknowledgments

W.W. is supported by the Royal Academy of Engineering (RF\201819\18\92). K.L.M. is supported by the Wellcome Trust (WT202788/Z/16/A). The Wellcome Centre for Integrative Neuroimaging is supported by core funding from the Wellcome Trust (203139/Z/16/Z).

## Supplementary Information

### Co-registration Details

Steps for co-registering the CAIPI-PF and conventional 3D multi-slab data with the MPRAGE image:

1. An intermediate b=0 image volume (denoted as *I*_1_) was acquired in the same scan as the MPRAGE image (with matched gradient distortions) using conventional 2D EPI and corrected for distortion using FSL’s “topup”^1, 2^ to minimize the impact of gradient nonlinearity distortions and improve the co-registration accuracy.
2. The b=0 images of CAIPI-PF and conventional 3D multi-slab sampling (denoted as *B*0_*CAIPI–PF*_ and *B*0_*conventional*_) were co-registered to *I*_1_ using FSL’s “flirt”^3, 4^ with default parameters to correct gradient nonlinearity distortions. The resultant images and transformations were denoted as *I*_2, *CAIPI–PF*_, *I*_2,*conventional*_ and *T*_2,*CAIPI–PF*,_ *T*_2_,_*conventional*_.
3. *I*_2,*CAIPI–PF*_ and *I*_2,*conventional*_ were co-registered to the MPRAGE image using FSL’s “epi_reg”^3, 4^. The resultant transformations were denoted as *T*_3,*CAIPI–PF*_ and *T*_3,*conventional*_.
4. *T*_2, *CAIPI–PF*_ and *T*_2,*conventional*_ were combined with *T*_3,*CAIPI–PF*_ and *T*_3,*conventional*_ using FSL’s “convert_xfm”^3, 4^. The resultant transformations were denoted as *T*_4, *CAIPI–PF*_ and *T*_4,*conventional*_.
5. *B*0_*CAIPI–PF*_ and *B*0_*conventional*_. were co-registered to the MPRAGE image using “flirt” with “bbr” as the cost function^5^ and *T*_4, *CAIPI–PF*_, *T*_4, *conventional*_ as initializations. The resultant transformations were used to resample the results from the diffusion analyses into the anatomical space.

**Figure S1.**
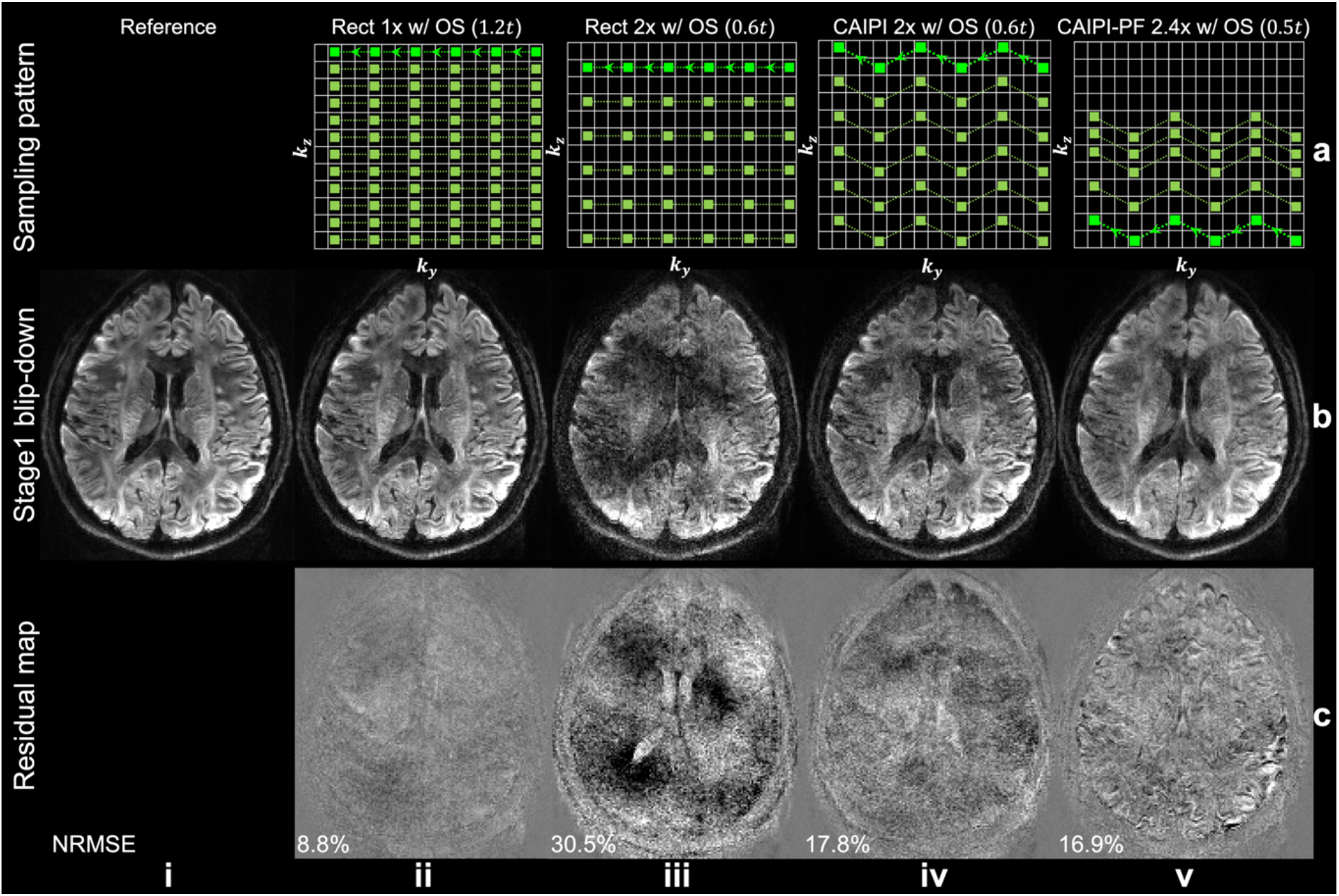
Stage 1 blip-down sampling and reconstruction. The sampling patterns (a), reconstruction results of a slab-central slice (b) from fully sampled reference (i) and different sampling patterns (ii-v) and their residuals with the fully sampled reference (c) for the blip-down data acquired with the evaluation protocol (1.22 mm isotropic resolution) are displayed, with relative scan times listed for each method. The trajectory of one shot of the multi-shot sampling is marked in bright green with arrows. The normalized root mean squared errors (NRMSE) of the whole slab are listed to quantify the image similarity. The parameter *t* represents the scan time of a rectangular sampling without over-sampling or acceleration along kz (as Fig. 2c, i).

**Figure S2.**
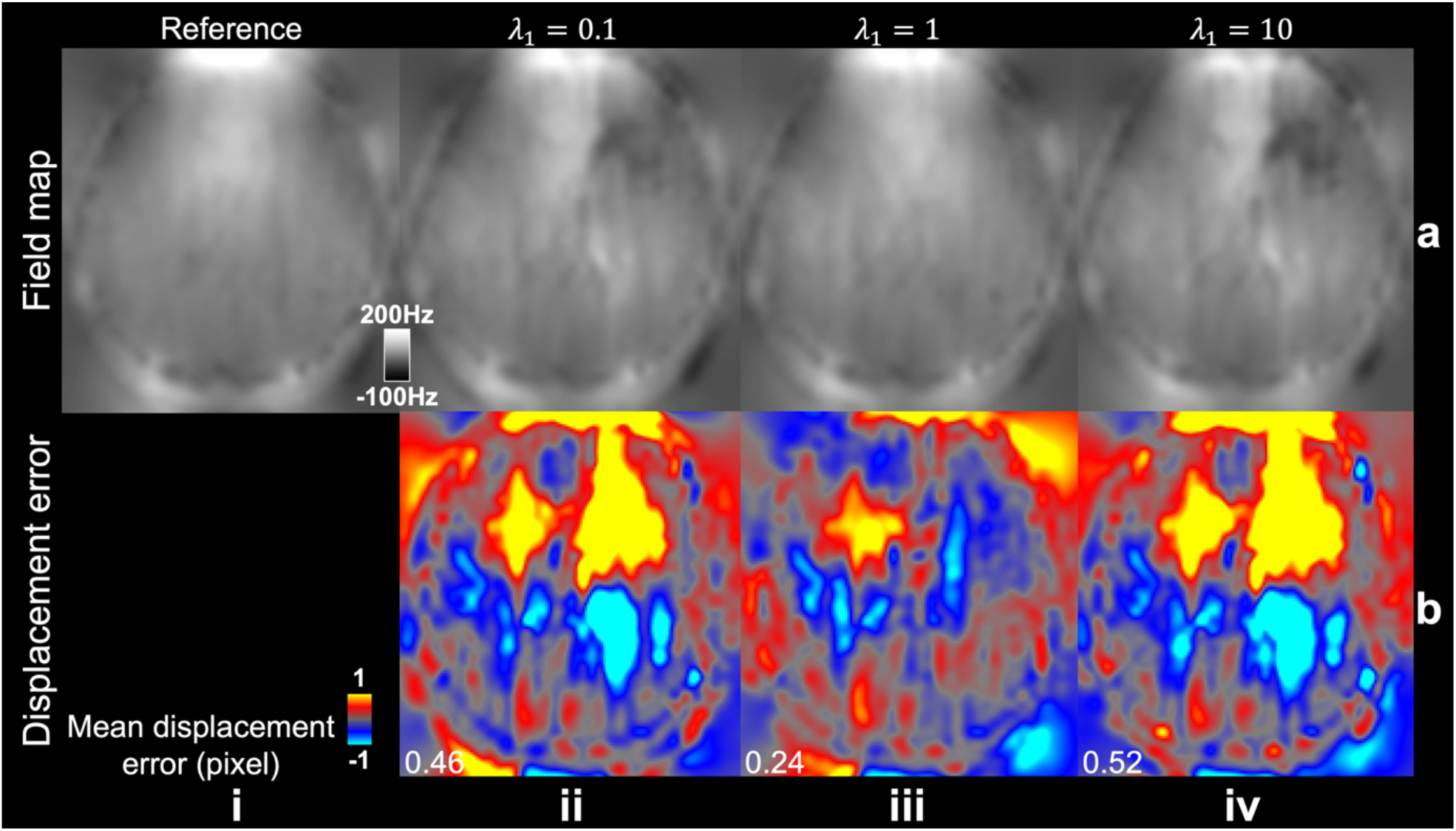
Impact of stage 1 SPIRiT weight on field map estimation. The field maps from “topup” estimated using reference blip-up/down data (a, i) and CAIPI–PF blip-up/down data with different SPIRiT weights for stage 1 reconstruction (a, ii-iv) and their voxel displacement error maps with the reference field map (b) acquired with the evaluation protocol (1.22 mm isotropic resolution) are displayed. The mean displacement errors of the whole slab are listed to quantify the image similarity.

**Figure S3.**
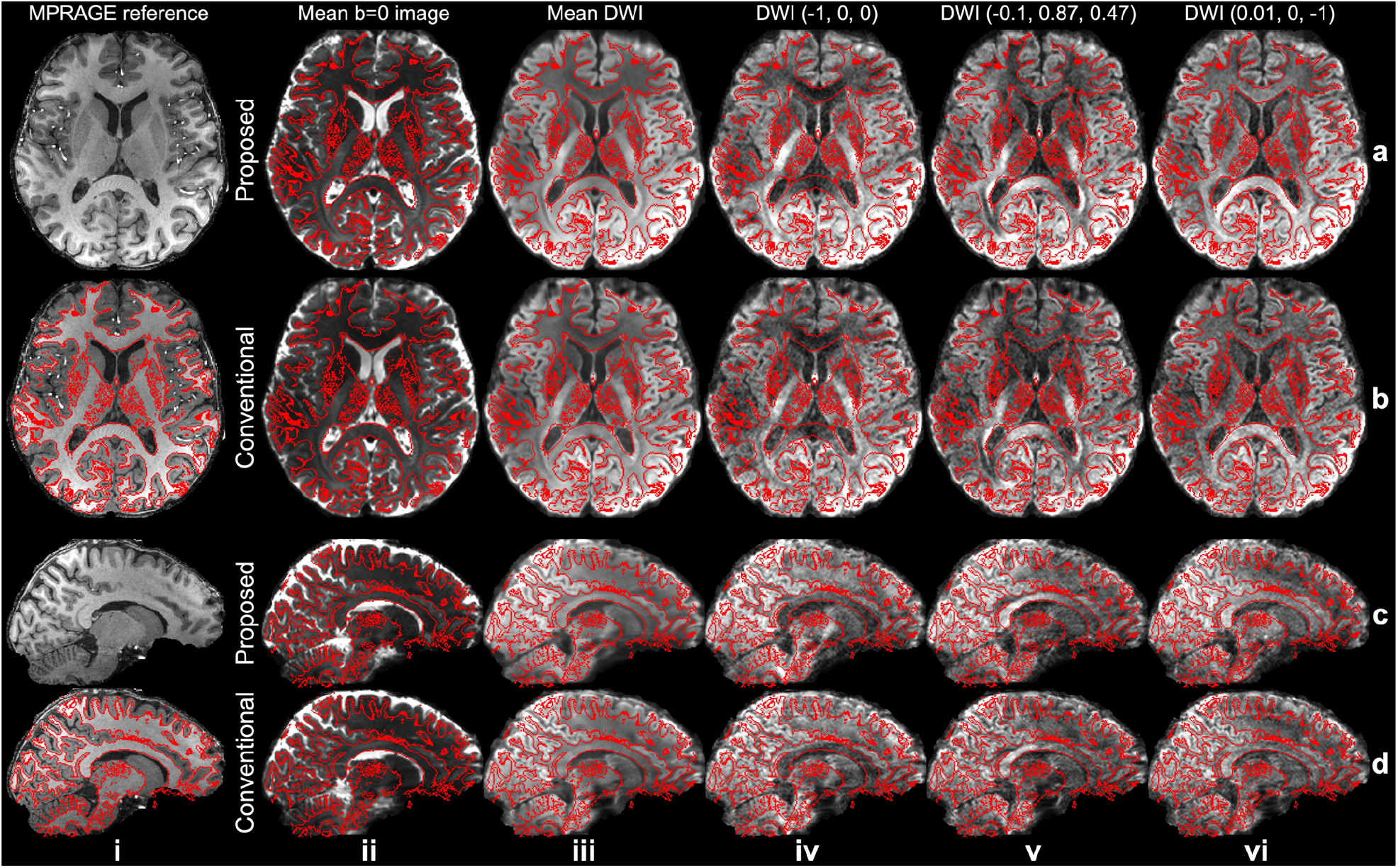
Comparison of image anatomical fidelity. The axial (a, b) and sagittal (c, d) views of the MPRAGE reference (0.86 mm isotropic resolution, i), mean b=0 images (ii), mean diffusion-weighted images (DWI) (iii), and three DWI along different diffusion directions (iv-vi) acquired with CAIPI–PF sampling (Proposed, a, c) and conventional 3D multi-slab sampling (Conventional, b, d) at 1.05 mm isotropic resolution from the same subject are displayed. The write matter boundary segmented by FSL’s “fast” is marked in red and overlayed. The diffusion data are co-registered to the MPRAGE reference for comparison.

**Figure S4.**
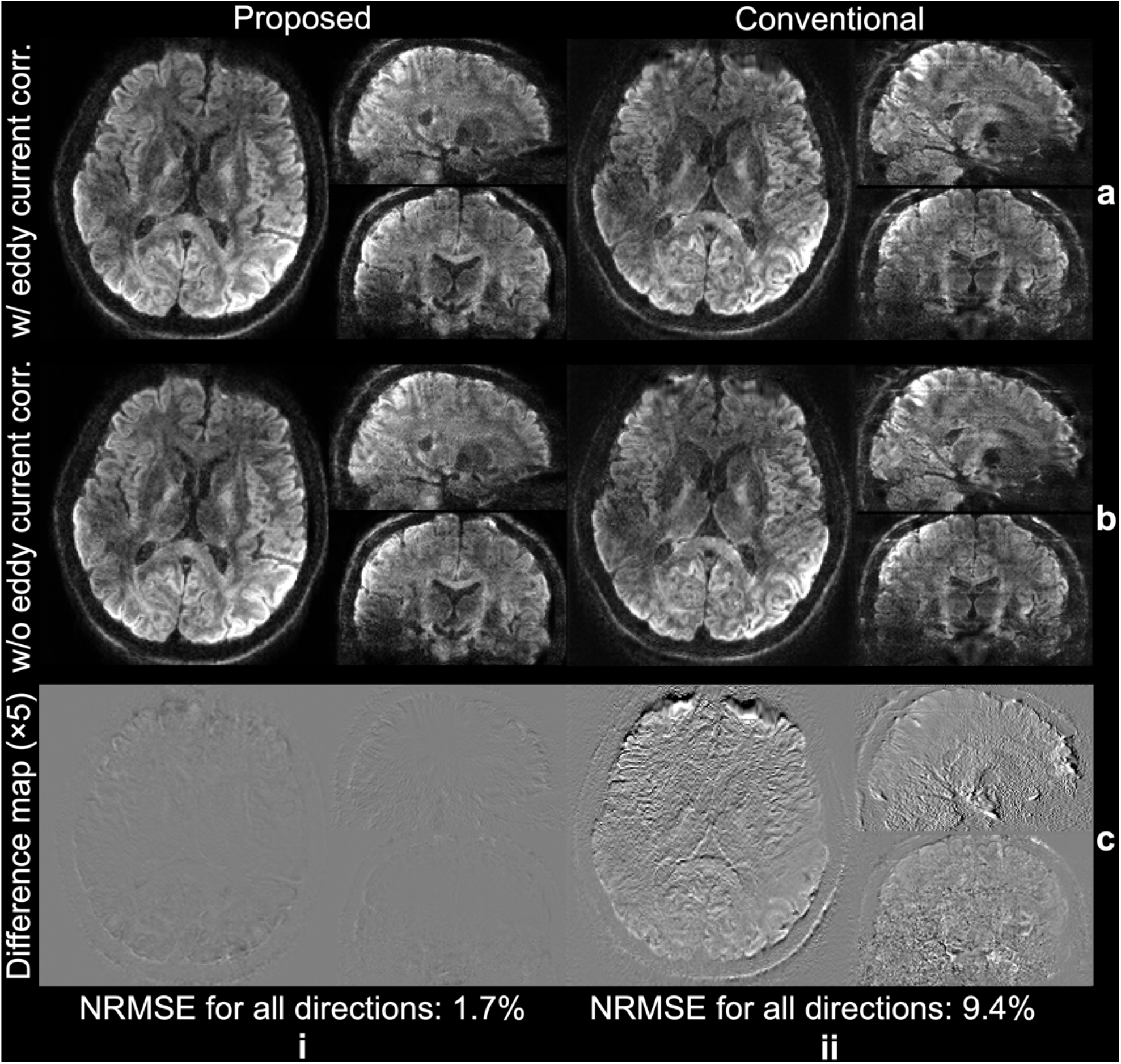
Correction of eddy current induced distortion. The single volume diffusion-weighted image along diffusion direction (−0.16, −0.63, 0.76) acquired with CAIPI–PF (Proposed, i) and conventional 3D multi-slab sampling (Conventional, ii) at 1.05 mm isotropic resolution processed by FSL’s “eddy” with eddy current correction (a), without eddy current correction (b), and their difference maps (c) of the same subject are displayed. The normalized root mean squared errors (NRMSE) for diffusion-weighted images of all diffusion directions are listed to quantify the image similarity.

**Figure S5.**
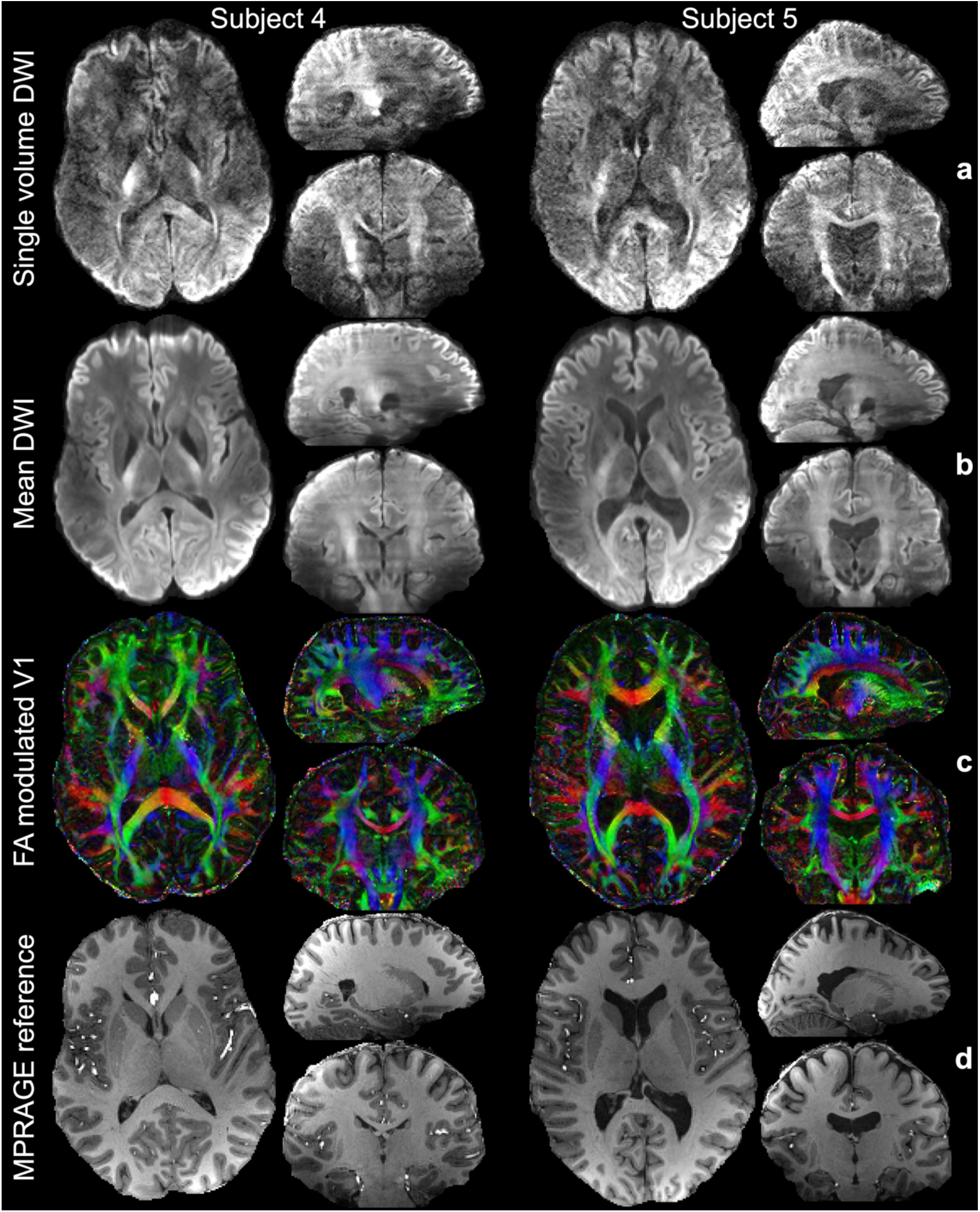
Diffusion image results of the remaining subjects. The single volume diffusion-weighted images (DWI) along (−0.26, −0.81, −0.52), the mean DWI (b), and the fractional anisotropy (FA) modulated primary eigenvector (V1) of diffusion tensor (c), and the anatomical images for reference (d) of two subjects are displayed. The diffusion images are acquired with CAIPI–PF sampling at 1.05 mm isotropic resolution. The anatomical images are acquired with MPRAGE at 0.86 mm iso. resolution and co-registered to diffusion images for comparison.

**Figure S6.**
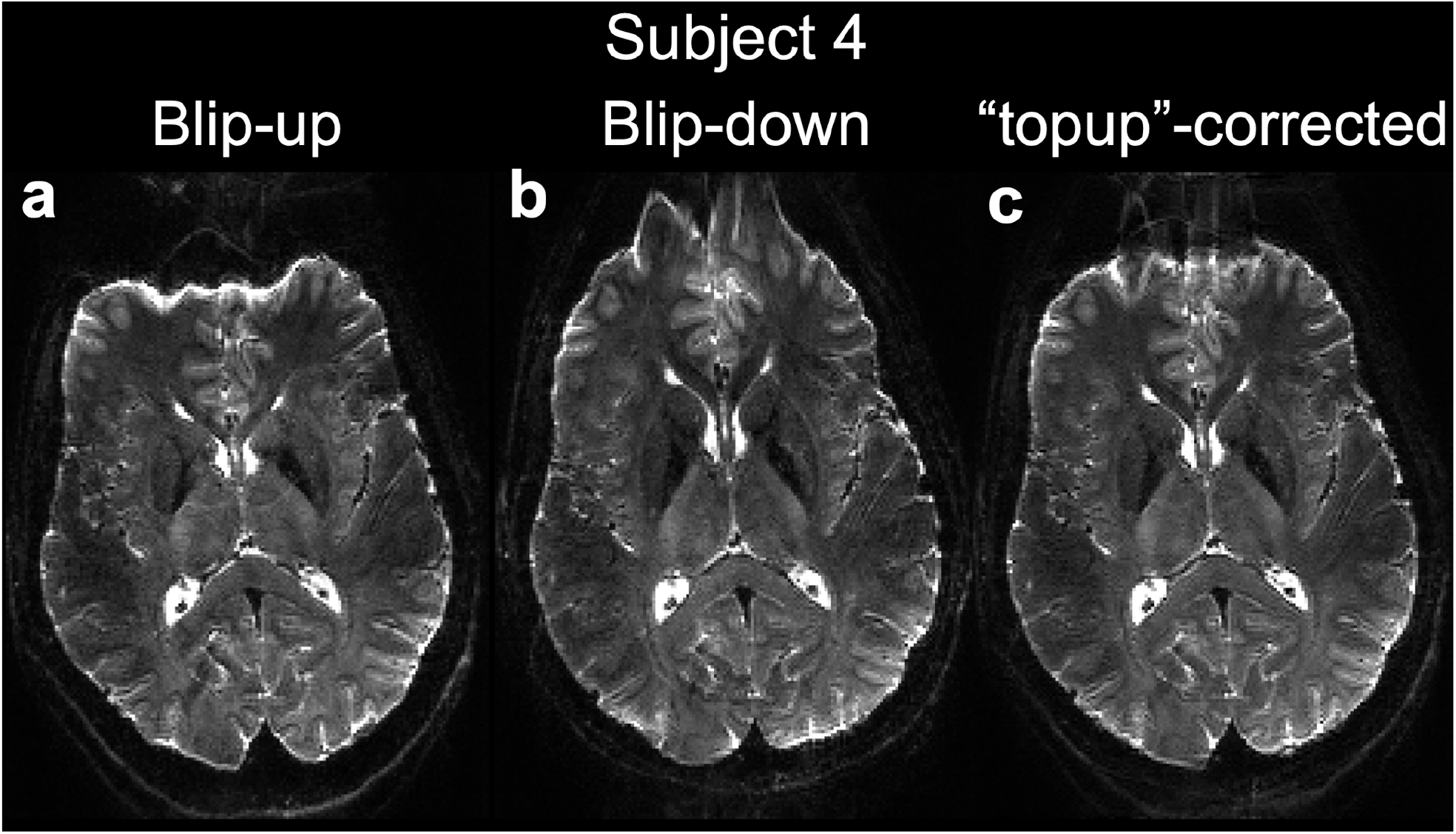
The b=0 images of a representative subject. An axial slice of blip-up (a), blip-down (b), and “topup”-corrected (c) b=0 images of a representative subject scanned with CAIPI–PF sampling at 1.05 mm isotropic resolution are displayed.

**Table S1.**
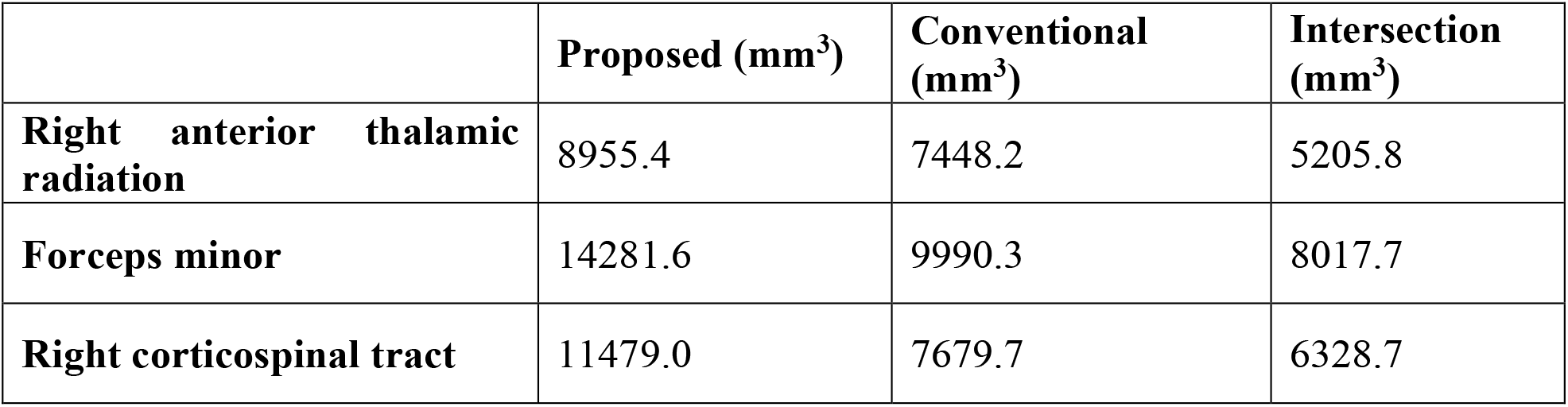
Tract mask volumes. Tractography results of data from proposed CAIPI–PF (Proposed) and conventional 3D multi-slab (Conventional) sampling are binarized to obtain tract masks for different tracts (threshold: 0.3%). Tract mask volumes are computed for tract masks of proposed and conventional data and their intersections.

**Figure S7.**
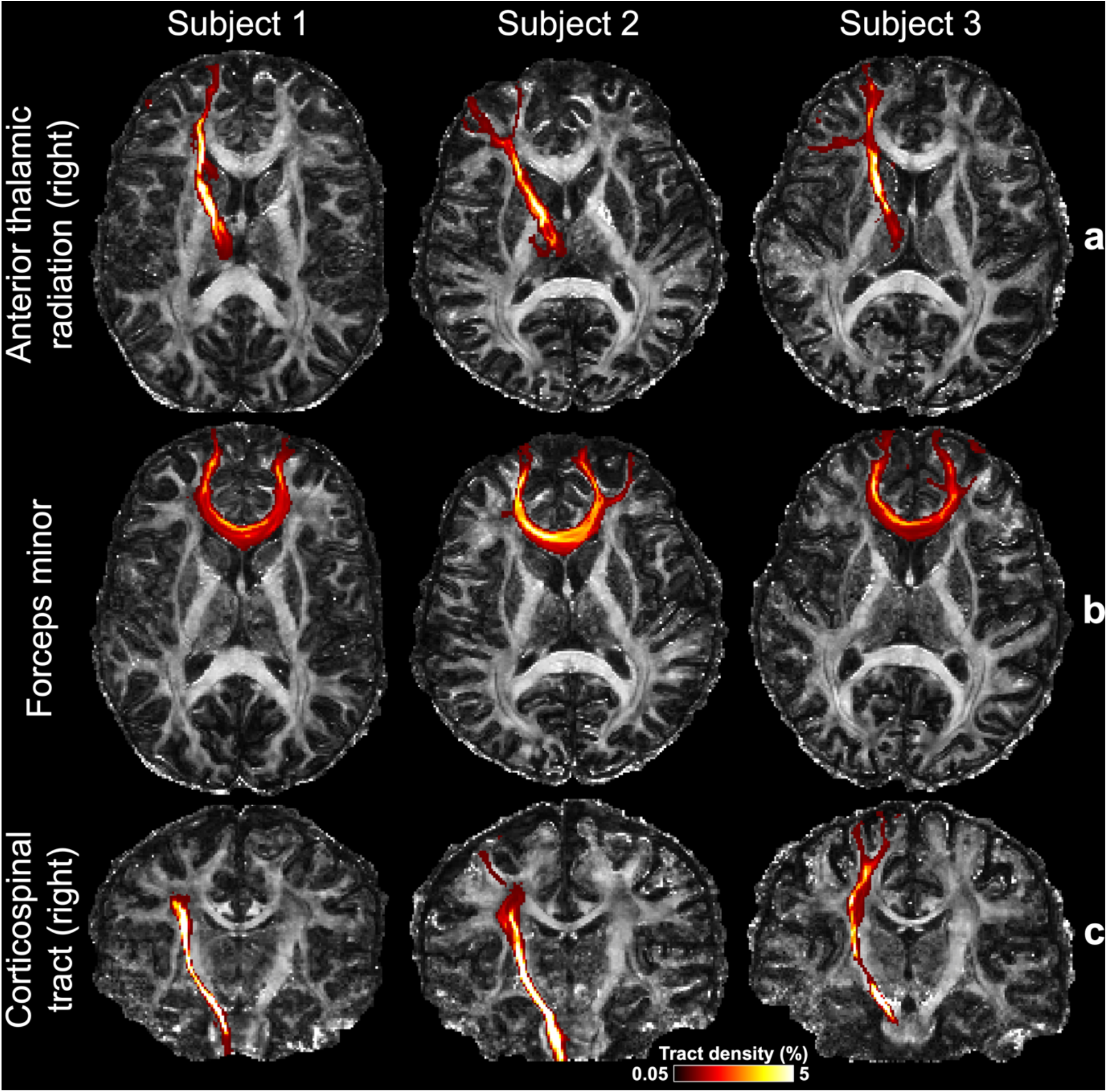
Tractography of multiple subjects. Tractography results (not maximum intensity projected) of three tracts including right anterior thalamic radiation (a), forceps minor (b), and right corticospinal tract (c) with tract density range 0.05%-5% from three subjects using the CAIPI–PF sampling (1.05 mm isotropic resolution) overlayed on their fractional anisotropy (FA) maps are displayed.

